# Suppression of Visceral Nociception by Selective C-Fiber Transmission Block Using Temporal Interference Sinusoidal Stimulation

**DOI:** 10.1101/2024.10.13.618090

**Authors:** Shaopeng Zhang, Longtu Chen, Eric Woon, Jia Liu, Jaehyeon Ryu, Hsin Chen, Hui Fang, Bin Feng

## Abstract

Chronic visceral pain management remains challenging due to limitations in selective targeting of C-fiber nociceptors. This study investigates temporal interference stimulation (TIS) on dorsal root ganglia (DRG) as a novel approach for selective C-fiber transmission block. We employed (1) GCaMP6 recordings in mouse whole DRG using a flexible, transparent microelectrode array for visualizing L6 DRG neuron activation, (2) ex vivo single-fiber recordings to assess sinusoidal stimulation effects on peripheral nerve axons, (3) in vivo behavioral assessment measuring visceromotor responses (VMR) to colorectal distension in mice, including a TNBS-induced visceral hypersensitivity model, and (4) immunohistological analysis to evaluate immediate immune responses in DRG following TIS. We demonstrated that TIS (2000 Hz and 2020 Hz carrier frequencies) enabled tunable activation of L6 DRG neurons, with the focal region adjustable by altering stimulation amplitude ratios. Low-frequency (20-50 Hz) sinusoidal stimulation effectively blocked C-fiber and Aδ-fiber transmission while sparing fast-conducting A-fibers, with 20 Hz showing highest efficacy. TIS of L6 DRG reversibly suppressed VMR to colorectal distension in both control and TNBS-induced visceral hypersensitive mice. The blocking effect was fine-tunable by adjusting interfering stimulus signal amplitude ratios. No apparent immediate immune responses were observed in DRG following hours-long TIS. In conclusion, TIS on lumbosacral DRG demonstrates promise as a selective, tunable approach for managing chronic visceral pain by effectively blocking C-fiber transmission. This technique addresses limitations of current neuromodulation methods and offers potential for more targeted relief in chronic visceral pain conditions.

## Introduction

Visceral pain is the primary symptom of disorders of gut-brain interactions (DGBIs) and a common reason for patients to seek gastroenterological care (Zia et al., 2022). Irritable bowel syndrome (IBS), a type of DGBIs, is estimated to affect up to 15% of persons in the USA (Lovell and Ford, 2012), with negative effects on quality of life and annual direct and indirect health care costs of ∼$30bn (Hulisz, 2004). Visceral pain is distinct from other types of pain, and current pharmacological treatments for IBS-related visceral pain are inadequate. Studies have shown that conventional pain medications (e.g., NSAIDs, acetaminophen, aspirin, and various opioids) do not significantly improve the symptoms of visceral pain (Chen et al., 2017a).

Neuromodulation, particularly electrical neurostimulation, has emerged as a potential non-pharmacological approach for managing chronic pain (Knotkova et al., 2021). Most clinical neurostimulators target the peripheral nervous system, including peripheral nerve field stimulation (Verrills et al., 2011), direct peripheral nerve stimulation (dPNS) (Günter et al., 2019), dorsal root ganglion (DRG) stimulation (Harrison et al., 2018), and spinal cord stimulation (SCS) (Linderoth and Foreman, 1999). SCS and dPNS are the most widely used neuromodulatory methods for chronic pain management. These techniques are based on the Gate Control Theory (Melzack and Wall, 1965), which suggests that activating large-diameter A-fibers can inhibit nociceptive transmission in the spinal cord dorsal horn. Activation of A-fibers by SCS or dPNS evokes a non-painful tingling sensation (paresthesia) that can “mask” pain sensation arising from the same area (Zhang et al., 2014). However, the Gate Control theory may not be fully applicable to treating visceral pain, as visceral organs have minimal innervation by A-fiber afferents (Feng and Gebhart, 2011;Feng and Guo, 2020). Also, conscious sensations arising from visceral afferent stimulation typically do not include tingling sensations associated with paresthesia (Jänig, 1996).

Unlike SCS and dPNS, DRG stimulation manages chronic pain in some patients without causing paresthesia (Verrills et al., 2019), suggesting additional underlying mechanisms beyond the Gate Control Theory. Using *ex vivo* single-fiber recordings, we pioneered in demonstrating that lumbosacral DRG stimulation in the sub-kilohertz range (20 to 100 Hz) reversibly blocks visceral afferent neural transmission in both control and visceral hypersensitive mice (Chen et al., 2019a;Chen et al., 2019b;Chen et al., 2022). Additionally, bilateral L6 DRG stimulation reversibly blocked the behavioral visceromotor response (VMR) to noxious colorectal distension (CRD), an objective assessment of visceral pain (Chen et al., 2022). The afferent blocking effect by sub-kilohertz DRG stimulation was later confirmed by studies using rat models of neuropathic pain (Chao et al., 2020;Chao et al., 2021), showing that low-frequency stimulation (20 Hz) blocked slow-conducting C-fiber afferents while leaving myelinated Aδ-fibers unblocked. Therefore, the pain-managing effect of DRG stimulation is likely contributed to by the transmission block of C-fiber afferents. Chronic visceral pain is characterized by maladaptive sensitization in both the peripheral and central nervous systems (Anand et al., 2007). Central sensitization is usually the outcome of prolonged peripheral sensitization, i.e., heightened peripheral input from sensitized afferents (Latremoliere and Woolf, 2009). In support, blocking sensitized afferent transmission with intracolonic lidocaine application effectively reduces visceral hypersensitivity in rodent models (Zhou et al., 2008) and relieves IBS-related pain in patients (Verne et al., 2003;Verne et al., 2005). Thus, DRG stimulation can be a promising strategy for ‘treating’ chronic visceral pain by suppressing the peripheral sensitization, i.e., selectively blocking the transmission of unmyelinated C-fiber visceral afferents.

Loss of therapeutic efficacy is the primary reason for the explantation of neurostimulator implants (Siegel et al., 2018). Enhanced programmability and tunability of the electrical fields delivered by implantable neurostimulators could potentially adapt to long-term changes at the electrode-tissue interface, thereby reducing the incidence of device explantation. We recently reported a novel design for a flexible array of 64 electrode contacts to interface with the L4 DRG in mice (Ryu et al., 2024), which couples with a programmable pulse generator (Biopro Scientific, LLC) to enable flexible delivery of electrical fields between any pair of electrode contacts (Liang et al., 2023). In addition to increasing the spatial distribution of electrode contacts, the tunability of the DRG neurostimulator can be further enhanced by implementing temporal interference stimulation (TIS), i.e., two kilohertz sinusoidal electrical stimulations of slightly different frequencies (Mirzakhalili et al., 2020). TIS has emerged as a non-invasive brain stimulation technique to modulate deep brain regions using surface-placed electrodes (Grossman et al., 2017). Recent studies have also applied TIS to stimulate motor neurons in the spinal cord (Sunshine et al., 2021). The key principles of TIS are: 1) Two or more oscillating electric fields are applied at a high ‘carrier frequency’ (e.g., 2000 Hz and 2020 Hz); 2) These fields interfere with each other, creating an envelope frequency at their difference, i.e., the ‘beat frequency’ (e.g., 20 Hz) deep within the brain or spinal cord; 3) This beat frequency envelope (e.g., 20 Hz sinusoidal wave) can modulate neuronal activity in targeted deep regions; 4) Neurons closer to the electrodes are comparatively less stimulated due to the high frequencies of the individual fields.

In this study, we aim to establish TIS in DRG stimulation for managing visceral pain arising from the distal colon and rectum. We assess the activation region of DRG neurons under TIS of different intensity ratios through optical GCaMP6 recordings. We also systematically study the nerve-blocking effects of sinusoidal stimulation using ex vivo single-fiber recordings. Furthermore, we conduct whole-animal behavioral assays to evaluate the effects of bilateral L6 DRG TIS in blocking the VMR to noxious CRD.

## Materials and Methods

All experiments were reviewed and approved by the University of Connecticut Institutional Animal Care and Use Committee. Mice were housed in pathogen-free facilities accredited by the American Association for Accreditation of Laboratory Animal Care and assured by the Public Health Service, following guidelines set forth in the Eighth Edition of the Guide for the Care and Use of Laboratory Animals. Housing consisted of individually ventilated polycarbonate cages (Animal Care System M.I.C.E.) with a maximum occupancy of 5 mice per cage. Environmental enrichment included nestlets and huts, with Envigo T7990 B.G. Irradiated Teklad Sani-Chips as bedding material. Mice were fed ad lib with either 2918 Irradiated Teklad Global 18% Rodent Diet or 7904 Irradiated S2335 Mouse Breeder Diet supplied by Envigo and supplied with reverse osmosis water chlorinated to 2 ppm using a water bottle. The animal housing facility maintained a 12:12 light-dark cycle, with ambient temperature regulated between 70-77°F (set point 73.5°F) and relative humidity between 35-65% (set point 50%). Animal care services staff performed daily health checks, and cages were changed biweekly.

### Implementing TIS using a flexible epidural electrode array and customized stimulator

As shown in the schematic in **Fig. 1A**, we delivered TIS to the DRG using two pairs of epidural surface electrodes. Each electrode pair delivers kilohertz sinusoidal current stimulation at 2000 and 2020 Hz, respectively. The temporal interference of the two sinusoidal sources results in an envelope frequency of 20 Hz at the focal region. This low-frequency component is capable of effectively modulating neural activity while the high-frequency carrier signals normally do not show obvious neural activation effects outside the focal region. Previous studies have demonstrated that the focal region can be shifted by adjusting the relative current amplitudes between the electrode pairs (Grossman et al., 2017;Mirzakhalili et al., 2020). To assess the tunability of the focal region, we delivered the same amount of current to the two pairs of electrodes with three different amplitude ratios: 1:3, 1:1, and 3:1.

**Figure 1.**
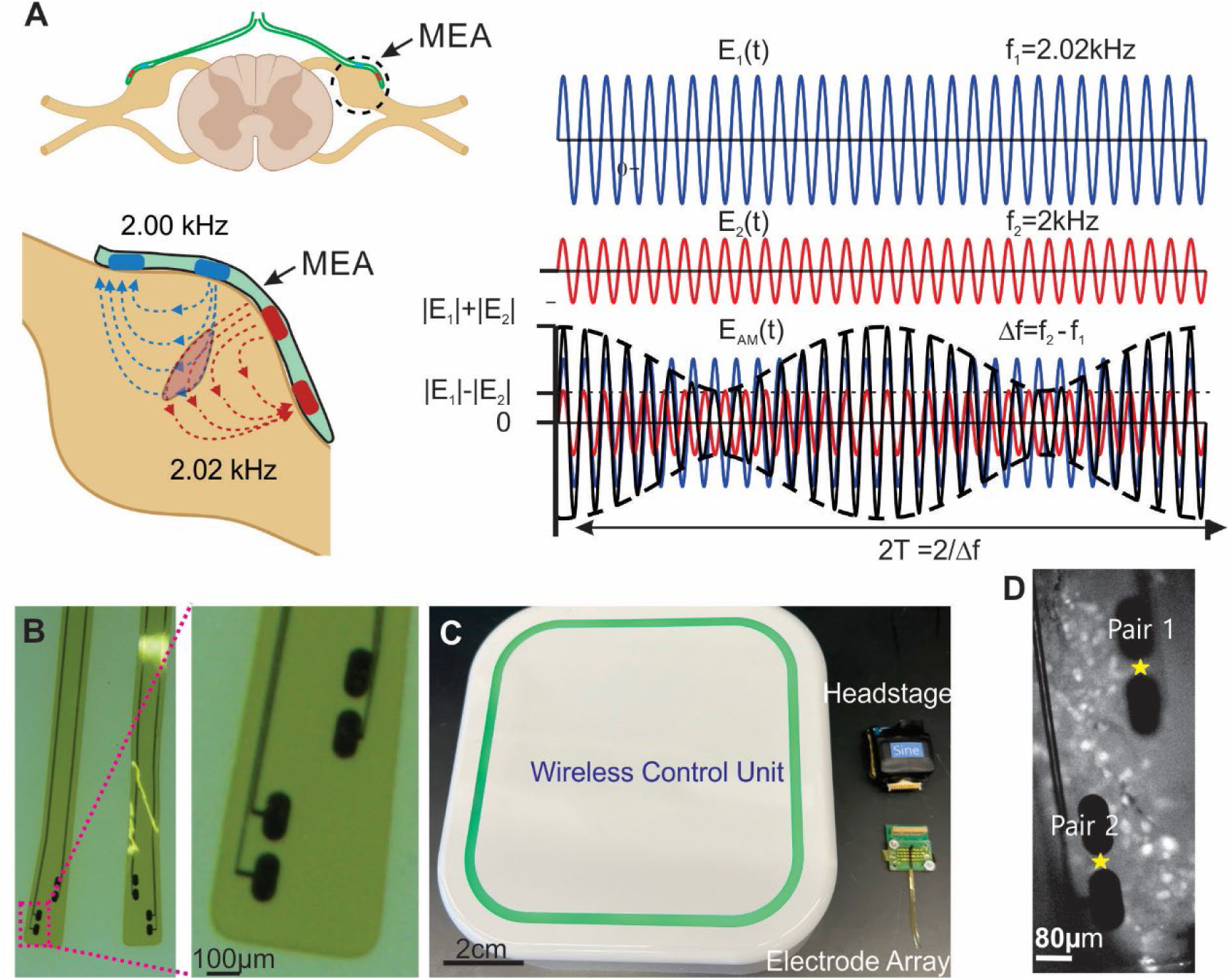
DRG stimulation by TIS via a flexible, transparent microelectrode array (MEA). (A) Schematic of TIS by two pairs of epidural surface electrodes on the DRG. (B) The MEA with two pairs of electrodes fabricated on a flexible and transparent Kapton/Polyimide substrate. It is designed to interface with L6 mouse DRG. Scale bar: 100 μm. (C) The custom-built wireless neural stimulation/recording system (Biopro Scientific, LLC.) consisting of a wireless control unit, headstage, and electrode array. (D) A representative image recording of GCaMP6s signals from a mouse L6 DRG. The TIS was delivered by the MEA that covers the dorsal surface of the L6 DRG. GCaMP6s signals were recorded through the transparent MEA.

As photographed in **Fig. 1B**, the stimulation microelectrode array (MEA) is fabricated on a soft substrate of Kapton/Polyimide (PI) to allow interfacing with bilateral mouse DRGs concurrently. The MEA is composed of six layers, including a Kapton/Polyimide (PI) bilayer substrate, Cr/Au/PEDOT:PSS stimulation electrodes with Cr/Au interconnections and a connection pad, and an SU-8 encapsulation layer. We designed two pairs of Au/PEDOT:PSS stimulation electrodes, each with a site area of 11,426.5 µm². PEDOT was electroplated to reduce impedance and enhance charge injection capacity (charge injection limit ±270 µA) (Dijk et al., 2021;Ryu et al., 2024). Compared to bare Au electrodes, PEDOT electrodes reduce irreversible electrochemical reactions, resulting in minimized pH fluctuations and the generation of reactive oxygen species. The device, with a thickness of approximately 9 µm, enables the MEA to conform to the contour surfaces of mouse DRGs as reported previously (Ryu et al., 2024), enhancing the reliability and effectiveness of electrical stimulation.

To deliver the TIS, we had the wireless neural stimulation/recording system customized by the manufacturer, i.e., Biopro Scientific, LLC., which stacked four off-the-shelf headstages (model # BPS-NL-035, Biopro Scientific, LLC) to achieve two-channel sinusoidal stimulations at 2000 and 2020 Hz, respectively (**Fig. 1C**). The hardware and firmware were reconfigured and reprogrammed to administer kilohertz sinusoidal current stimulation across a broad frequency range (20 to 2020 Hz). To ensure charge-balanced stimulation and prevent tissue damage, a 0.1 µF capacitor was incorporated in series with the stimulus pathway. The BPS system features a web browser-based software interface, enabling wireless adjustment of stimulus parameters during experiments and providing flexible control over neuromodulation protocols.

### Assessing the activation of DRG neurons by TIS via GCaMP6 Imaging

To systematically determine the spatial precision and tunability of DRG stimulation by TIS, we conducted ex vivo high-throughput calcium imaging on whole DRG harvested from transgenic mice as reported in our prior studies (Bian et al., 2021;Guo et al., 2021). L6 DRGs were harvested from mice heterozygous for GCaMP6s-LoxP and VGLUT2-Cre genes and interfaced with the flexible microelectrode array (MEA) as shown in **Fig. 1B**. GCaMP6s calcium transients were recorded through the transparent MEA when TIS was delivered to the DRG through the two pairs of surface electrodes as shown in **Fig. 1D**. The aforementioned TIS stimulation protocols with the same total current amplitude but different ratios between pairs were evaluated (1:3, 1:1, 3:1). GCaMP6s signals were recorded at 960 by 540 pixels and 30 frames per second through the transparent MEA using a high-speed, ultra-low noise sCMOS camera (Xyla-4.2P, 82% quantum efficiency, Andor Technology, CT). To enhance the contrast of images and facilitate observation of the activated neurons, we developed a customized MATLAB program to reveal regions with increased brightness by applying a median filter and subtracting the background image. For each stimulation protocol, we identified activated neurons and calculated the centroid of their positions. The distances (D1 and D2) from this centroid to the centers of electrode Pairs 1 and 2 were measured and normalized to the inter-pair distance.

### Assess the effect of sinusoidal neuromodulation through ex vivo single-fiber recordings

We systematically assessed the effect of low-frequency sinusoidal current stimulation on peripheral nerve transmission via single-fiber recordings from mouse sciatic nerve axons. Sciatic nerves were harvested from adult C57BL/6 mice of both sexes (10-16 weeks, 25-35 g) following a previously reported surgical procedure (Chen et al., 2017b;Zhang et al., 2024). Briefly, mice were anesthetized with 2% isoflurane inhalation and euthanized by transcardiac perfusion with oxygenated Krebs solution (in mM: 117.9 NaCl, 4.7 KCl, 25 NaHCO_3_, 1.3 NaH_2_PO_4_, 1.2 MgSO_4_, 2.5 CaCl_2_, and 11.1 D-glucose at room temperature, bubbled with 5% CO_2_ and 95% O_2_). The perfused carcass was immediately transferred to a dissection chamber circulated with oxygenated ice-cold Krebs solution for nerve harvesting.

As illustrated in **Fig. 2A**, harvested nerves were transferred to a custom-built two-compartment chamber consisting of a Krebs chamber and a mineral oil chamber (Chen et al., 2017b). The proximal end of the nerve was pinned in the Krebs chamber, which was perfused with oxygenated Krebs solution at 28-30°C. The distal end (∼5 mm) was gently pulled through a custom-built gate to the mineral oil chamber for extracellular single-fiber recordings. The distal ∼10 mm end of the nerve in the oil chamber was split into fine filaments (∼10 µm thickness) to enable recordings of action potentials (APs) from individual axons. A customized 5-channel electrode array (Pt/Ir, A-M System) was used to interface with these split nerve filaments. Single-unit APs were recorded simultaneously from all five channels, digitized at 15 to 20 kHz, and band-pass filtered (200-3000 Hz) using an Intan RHD USB interface board. The multichannel recording signals were also monitored by a CED 1401plus data acquisition system and stored on a PC using Spike2 software (v7.1, CED, Cambridge, UK).

**Figure 2.**
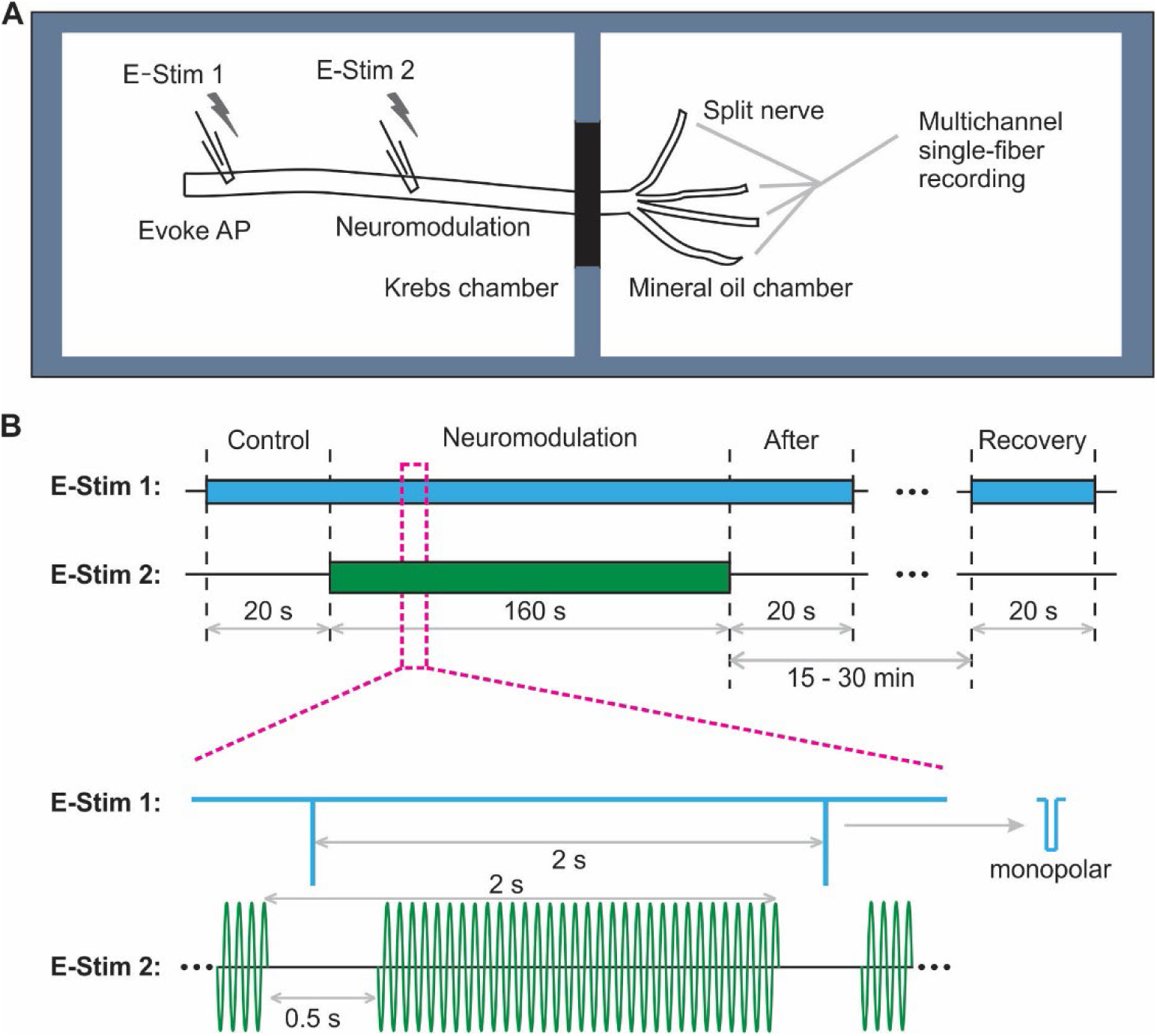
Experimental setup and protocol for assessing axonal transmission block by sinusoidal neuromodulation. (A) Schematic of the ex vivo two-chamber setup for single-fiber recordings from peripheral nerves. The Krebs chamber houses the proximal nerve end and stimulation sites (E-Stim 1 and E-Stim 2), while the mineral oil chamber enables multichannel single-fiber recordings from split nerve endings. (B) Synchronized stimulation protocol illustrating the timing of evoked action potentials (E-Stim 1, blue) and neuromodulation (E-Stim 2, green). The protocol consists of Control, Neuromodulation, After, and Recovery phases. The expanded view shows the temporal relationship between monopolar pulse stimulation at E-Stim 1 and sinusoidal stimulation trains at E-Stim 2.

As illustrated in **Fig. 2B**, we implemented a synchronized stimulation protocol to characterize the instantaneous effect of sinusoidal neuromodulation on axonal AP transmission. Neural transmission was initiated via 0.5 Hz pulse stimulation at the proximal end of the harvested nerve, indicated by “E-Stim 1” in **Fig. 2A** (cathodic constant current, 0.2 ms pulse width) from a programmable stimulation isolator (Model 4100, A-M Systems, WA). To modulate the evoked AP transmission, sinusoidal stimulation at 20 to 2000 Hz was delivered at the site marked as “E-Stim 2” in **Fig. 2A** using a separate programmable stimulation isolator (Model 4100, A-M Systems, WA). Electrical stimulation at both stimulation sites was delivered through custom-built suction electrodes by pulling quartz glass capillaries with tips approximately 30% smaller than the diameter of the sciatic nerve trunk (∼Φ400 µm) (Zhang et al., 2024).

The synchronized stimulation protocol consisted of four phases: Control (20 s), Neuromodulation (160 s), Immediately After Neuromodulation (20 s), and Recovery (20 s, 15-30 min after terminating the neuromodulation). During the Neuromodulation phase, E-Stim 1 delivered monopolar pulses every 2 s to evoke APs, while E-Stim 2 applied continuous trains of sinusoidal stimulation for 1.5 s followed by a 0.5 s inter-train interval. This synchronized protocol allowed for instantaneous monitoring of subtle changes in AP transmission during sinusoidal neuromodulation, with the 0.5 s interval between neuromodulation trains providing unobstructed observation of evoked APs. Stimulus thresholds were determined at both stimulation sites, with E-Stim 1 set to exceed 5 times the threshold amplitude (0.1-1.5 mA) for robust AP generation. Neuromodulation intensities at E-Stim 2 were set as suprathreshold, specifically at ∼150% of the threshold determined for 20 Hz stimulation. This suprathreshold stimulation was essential for achieving effective conduction block, as subthreshold stimulation has been shown to be ineffective in blocking neural transmission in our previous studies (Chen et al., 2022;Zhang et al., 2024).

### The effect of TIS of L6 DRG in suppressing visceral nociception

To quantitatively determine the suppression of visceral nociception by TIS of L6 DRG, we conducted in vivo behavioral experiments to measure the visceromotor responses (VMR) to graded colorectal distension (CRD) by an intracolonic balloon in anesthetized mice (**Fig. 3A**). The VMR was quantified by electromyographic (EMG) recordings from abdominal oblique musculature before and after TIS. C57BL/6 mice of both sexes (8-12 weeks, 25-35 g) were used for all experiments. Mice were randomly assigned to two groups: the TNBS group and the saline group. The TNBS group received intracolonic administration of 0.2 ml 2,4,6-trinitrobenzenesulfonic acid (TNBS) solution (10 mg/ml in 50% ethanol; Sigma-Aldrich, St. Louis, MO, United States) via a 22-gauge feeding needle (#18061-22, Fine Science Tools, CA) to induce visceral hypersensitivity following our previous report (Feng et al., 2012b). The control group received intracolonic treatment of an equivalent volume of saline. VMR measurements were conducted 10-14 days post-treatment, a period previously characterized to show significant behavioral visceral hypersensitivity in TNBS-treated mice (Feng et al., 2012b).

**Figure 3.**
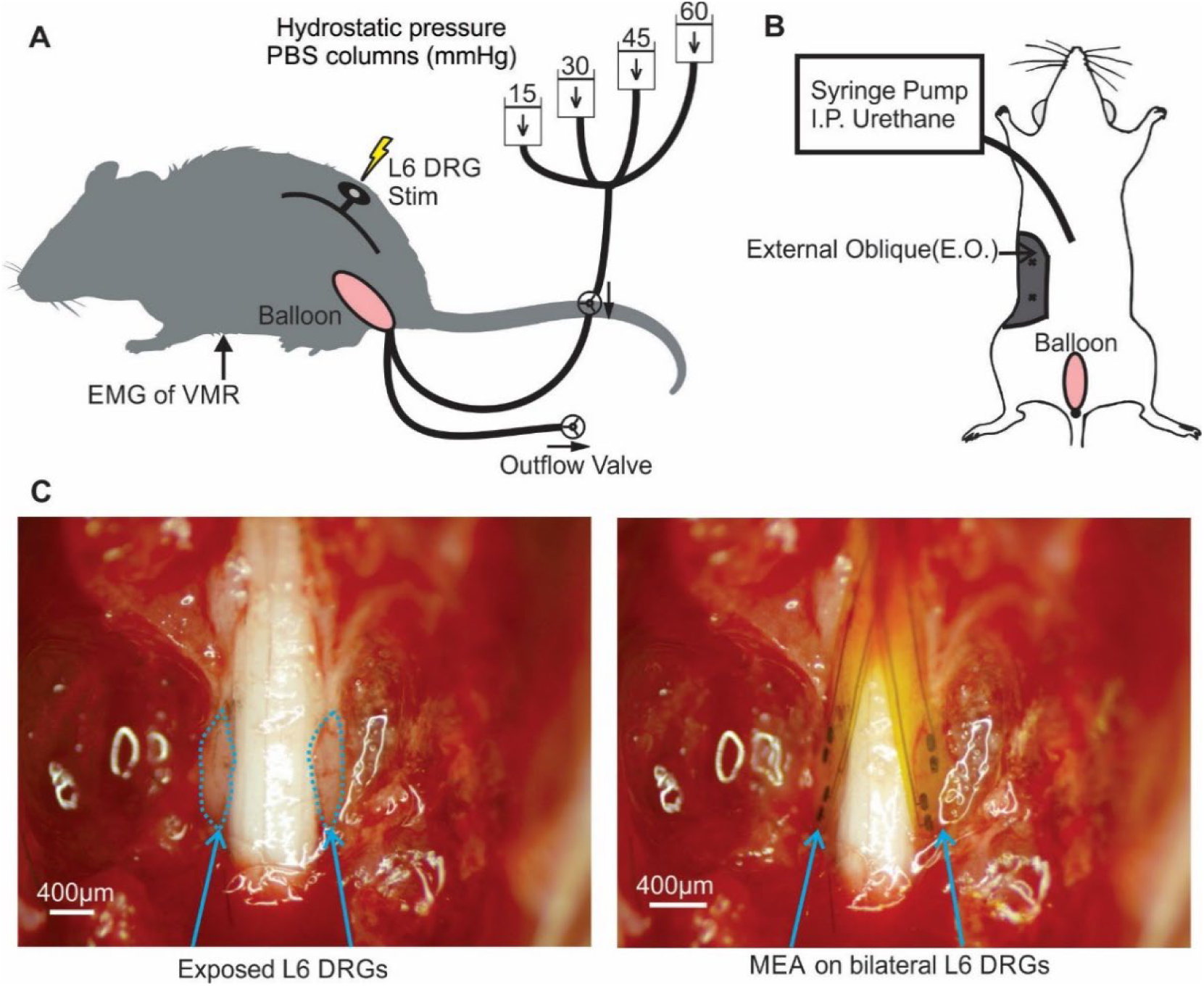
Experimental setup for measuring visceromotor responses (VMR) to colorectal distension (CRD) in mice. (A) Schematic of the in vivo preparation showing L6 DRG stimulation, colorectal balloon distension, and EMG recording of VMR. Hydrostatic pressure columns (15, 30, 45, and 60 mmHg) connected to the balloon enable graded CRD. (B) Ventral view illustrating electrode placement on the external oblique (E.O.) muscle for EMG recording and the intraperitoneal (I.P.) catheter for continuous urethane infusion via syringe pump. (C) In vivo application of the MEA on bilateral L6 DRGs. Left: Exposed L6 DRGs. Right: MEA placement on bilateral L6 DRGs. Scale bar: 400 μm.

The surgery procedures and hardware setup for the in vivo experiment have been reported in detail in our previous publication (Zhang et al., 2023). Briefly, on the day of a VMR experiment, mice were anesthetized with 2% isoflurane inhalation and placed on a feedback-controlled heating pad (HP-150, Auber Instruments, Alpharetta, GA, United States) to maintain body temperature at ∼37°C (Zhang et al., 2023). During the surgery, the anesthetized mouse was first placed in a supine position. Teflon-coated stainless steel wire electrodes (Conner Wire, Chatworth, CA, United States) were used to record abdominal EMG activity. The electrode tips (∼0.5 mm exposed) were sutured directly to the external oblique (E.O.) musculature immediately above the inguinal ligament using hypoallergenic bioabsorbable sutures (polyglactin, Ethicon), with 1 mm separation between electrodes (**Fig. 3B**). EMG signals were amplified using a differential amplifier (Model 1700, A-M Systems, Sequim, WA, United States) and digitized using a CED 1401 interface (Cambridge Electronic Design Limited, Cambridge, United Kingdom). A small abdominal incision was made for a thin polyethylene catheter (Φ 0.2 mm) to be placed intraperitoneally for continuous low-dose urethane infusion via a syringe pump throughout the experiment (**Fig. 3B**). Then, the mouse was placed in a prone position for the surgery to expose bilateral L6 DRG. The dorsal process and pedicles of the L6 vertebrae were carefully removed using a high-speed dental drill (Chen et al., 2022). Customized MEAs shown in **Fig. 1B** were placed on bilateral L6 DRGs for delivering the TIS. The left panel in **Fig. 3C** shows the exposed L6 DRGs, while the right panel demonstrates the placement of the MEA over the DRGs. Afterwards, a custom-made thin-film polyethylene balloon (Φ mm × 15 mm when distended) was inserted intra-anally into the colorectum. The balloon depth was precisely controlled to be 5-10 mm from the anus by measuring the distance between the distal end of the balloon and the anus (Zhang et al., 2023). The balloon was connected to a custom-built distension device consisting of four hydrostatic water columns set at 15, 30, 45, and 60 mmHg pressures, respectively. The onset and termination of CRD were regulated by computer-controlled solenoid valves.

After completing the surgical procedures, mice were transitioned from isoflurane to urethane anesthesia for hours-long recording of VMR to CRD following our recently established anesthesia protocol (Zhang et al., 2023). Briefly, an initial dose of 0.6 g/kg urethane was administered via the i.p. catheter, and isoflurane was concurrently turned off. Following a 60-min period of isoflurane clearance, mice were given a continuous i.p. infusion of low-dose urethane [0.15–0.23 g per kg weight per hour (g/kg/h)] through the implanted catheter using a syringe pump (World Precision Instruments, Sarasota FL). This in vivo setup enabled reliable assessment of visceromotor responses (VMR) to noxious colon distension (CRD) under urethane anesthesia, providing a reliable baseline for assessing the effect of L6 DRG stimulation.

### Immunohistological staining to assess the immediate immune responses in the DRG

We conducted histological staining of the targeted DRG to evaluate the possible immunoreaction caused by kilohertz DRG stimulation. L6 DRGs undergoing hours-long TIS electrical neuromodulations were harvested after each experiment and fixed in 10% buffered formalin overnight at 4°C. The adjacent L5 DRGs from the same mouse that did not receive direct stimulation were also collected as controls. The fixed tissue was washed with phosphate-buffered saline (PBS) three times (10 min each) to remove the fixative, cryoprotected in 20% sucrose in PBS, embedded in OCT compound (Sakura Finetek, Tokyo, Japan), frozen, and sectioned at 20 µm thick. Macrophages were stained using rat anti-F4/80 (1:500, Abcam, Cambridge, MA) and Cy3-conjugated anti-rat IgG (1:250), and nuclei were stained with DAPI. Fluorescent images were captured with a 60x oil objective (60X/1.40 Plan Apo oil, DIC, Nikon) using a Nikon A1R confocal microscope with the z-step set at 0.5 µm. Detection of the fluorescent signal threshold was reported previously (Guo et al., 2019). Using ImageJ, we projected each image stack from the myenteric plexus with “maximum intensity *z*-projection” to convert into an 8-bit grayscale image. The threshold was set to exclude at least 99.99% of the background intensity based on its Gaussian distribution. The threshold value was calculated from the standard normal distribution (Z) equation: threshold=3.72 SD + M, where 3.72 was the *Z*-score at *P* = 0.9999, SD and M were the standard deviation and mean of background intensities in 8-bit grayscale (0-255), respectively. The area fractions of fluorescent pixels corresponding to F4/80 and DAPI were separately calculated using ImageJ, and the ratio of the F4/80 area fraction to the DAPI area fraction was used to quantify the relative intensity of F4/80.

## Data Analysis and Statistics

Action potentials recorded through single-fiber recordings were processed off-line using custom MATLAB scripts. The detection threshold for individual action potentials was set at four times the root mean square (RMS) amplitude of the background noise, measured from a 10 ms window preceding each stimulation. EMG activities evoked by CRD were recorded from the abdominal oblique musculature, digitized at 2000 Hz, and processed off-line using customized MATLAB scripts. The EMG signals were rectified for calculating the area under the curve (AUC), which was used to evaluate the level of VMR to CRD. VMR evoked by CRD was quantified as the AUC values during the 5 s CRD subtracted by the AUC of the 5 s pre-distending baseline recording. Data are presented as means ± standard error of the mean (SEM). Statistical analyses were performed using SigmaStat v4.0 (Systat Software, San Jose, CA). ANOVA was used for comparisons between multiple groups, followed by Bonferroni post-hoc comparisons. Differences were considered statistically significant when p < 0.05.

## Results

### TIS delivered by epidural surface MEA enables tunable spatial activation of L6 DRG neurons

We used GCaMP6 recordings as a key metric to assess the activation of afferent neurons in the L6 DRG by TIS. Representative GCaMP6s recordings from the intact L6 DRG, captured through the transparent MEA, are shown in **Fig. 4A**. Typical GCaMP6s signals before and after TIS are presented in **Figs. 4A1** (baseline) and **4A2** (post-stimulation), respectively. **Figs. 4A3** to **4A7** highlight neurons activated by both interference and non-interference stimulation configurations, with neurons showing a significant increase in GCaMP6f signal identified by subtracting the baseline signal. In **Figs. 4A3** and **4A4**, identical stimulation frequencies were applied to both electrode pairs (non-interference) at 2000 Hz and 20 Hz, respectively. In **Figs. 4A5** to **4A7**, TIS was delivered at 2000 Hz and 2020 Hz, with amplitude ratios for electrode pair 1 versus pair 2 set to 1:1, 1:3, and 3:1, respectively. Under identical stimulation amplitudes, non-interference kilohertz stimulation (**Fig. 4A3**) elicited minimal neuronal activation, while sub-kilohertz stimulation (**Fig. 4A4**) triggered non-selective activation across most DRG areas. In contrast, TIS selectively activated a subset of DRG neurons (**Fig. 4A5**). **Fig. 4B** shows the normalized fluorescence intensity of two representative neurons (indicated by arrows in **Fig. 4A2**) over time, demonstrating a clear onset of activation in response to TIS.

**Figure 4.**
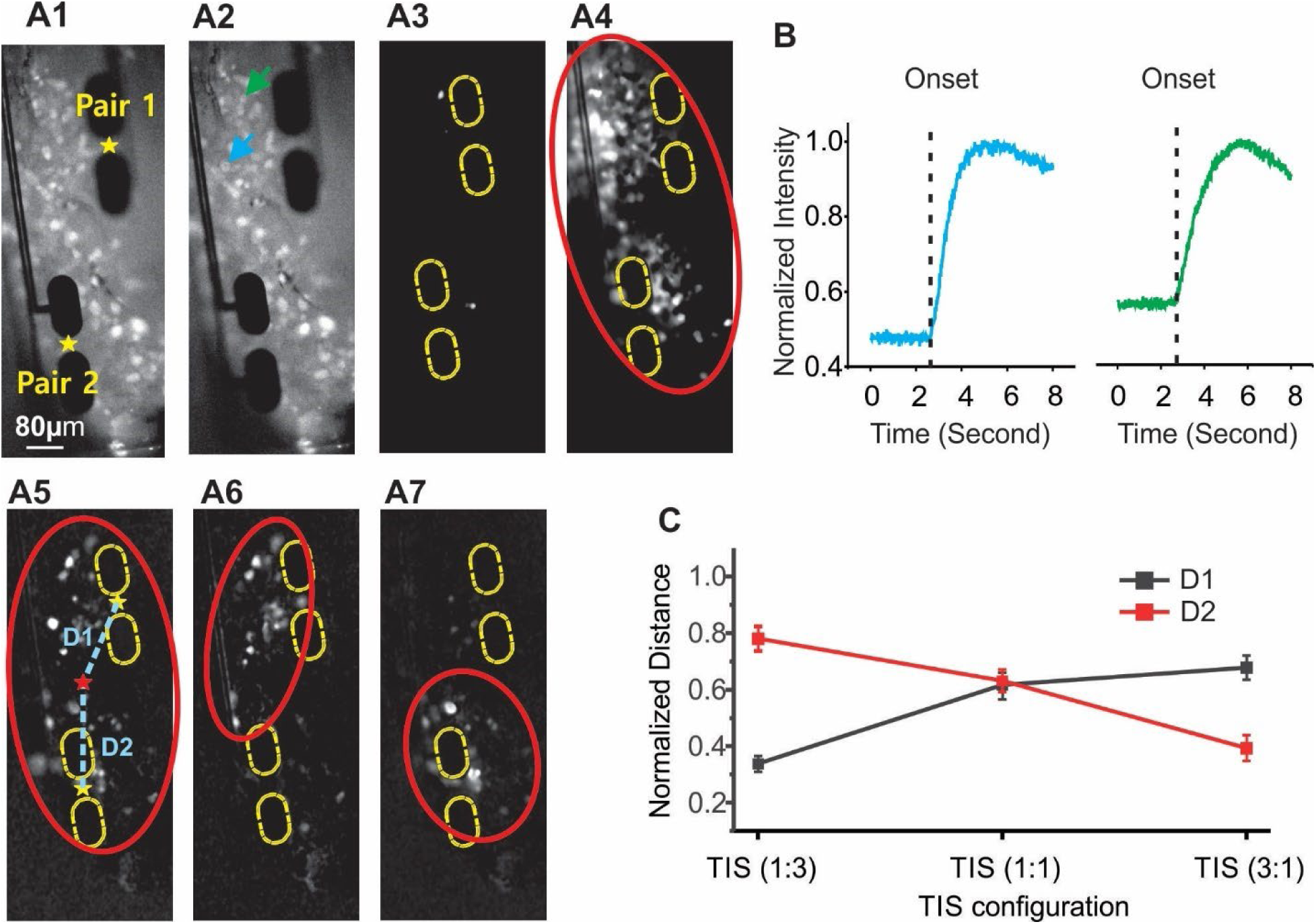
Spatial Control of Neuronal Activation by Temporal Interference Stimulation (TIS). (A1–A7) Representative fluorescent GCaMP6s recordings from the whole L6 DRG under different TIS configurations, with amplitude ratios of 1:1, 1:3, and 3:1. (B) Normalized fluorescence intensity over time from two activated DRG neurons, indicated by arrows in A2. (C) Distances (D1 and D2) between the centroid of activated neurons (encircled by an ellipse) and each of the two electrode pairs. Distances were normalized by the distance between the electrode pairs. Error bars represent the standard error of the mean (n = 8).

Activated DRG neurons were quantified by applying a threshold that excluded at least 99.99% of the background intensity (see § Methods). An ellipse was drawn to encircle over 95% of the activated neurons. The distances between the ellipse’s centroid and electrode pairs 1 and 2 were measured as D1 and D2, respectively. These distances, measured from eight DRGs, were normalized by the distance between electrode pair 1 and 2 and are shown in **Fig. 4C**. D1 differed significantly across the three TIS configurations (1:3, 1:1, and 3:1) based on one-way ANOVA with repeated measures (p < 0.05 for all post-hoc comparisons), and the same was observed for D2 (p < 0.05 for all post-hoc comparisons). Additionally, D1 and D2 were significantly different in the 1:3 (paired t-test, p < 0.01) and 3:1 (p < 0.01) TIS configurations. However, for the 1:1 ratio, D1 and D2 were comparable (p = 0.91).

### Characterization of nerve activation by sinusoidal current stimulation with single-fiber recordings

To evaluate the efficacy of sinusoidal stimulation compared to conventional pulse stimulation, we systematically assessed activation thresholds across a range of frequencies using single-fiber recordings from split peripheral nerve axons in the ex vivo preparation described in **Fig. 2**. Stimulation was applied at the E-Stim 2 site, and activation thresholds were defined as the minimum stimulating amplitude required to reliably evoke action potentials. **Figs. 5A** and **5B** show representative single-fiber recordings from individual nerve axons activated by electrical pulse stimulation (biphasic, cathodic-first, 200 μSec pulse width) and sinusoidal stimulation (20 Hz), respectively. Pulse stimulation allows precise determination of axonal conduction velocity based on the conduction delay. Spikes evoked by sinusoidal stimulation can be identified by applying band-pass filtering to remove stimulus artifacts. For each of the 10 C-fiber axons tested, we systematically determined activation thresholds for pulse stimulation (200 μSec) and six sinusoidal stimulations with frequencies ranging from 20 Hz to 2 kHz.

**Figure 5.**
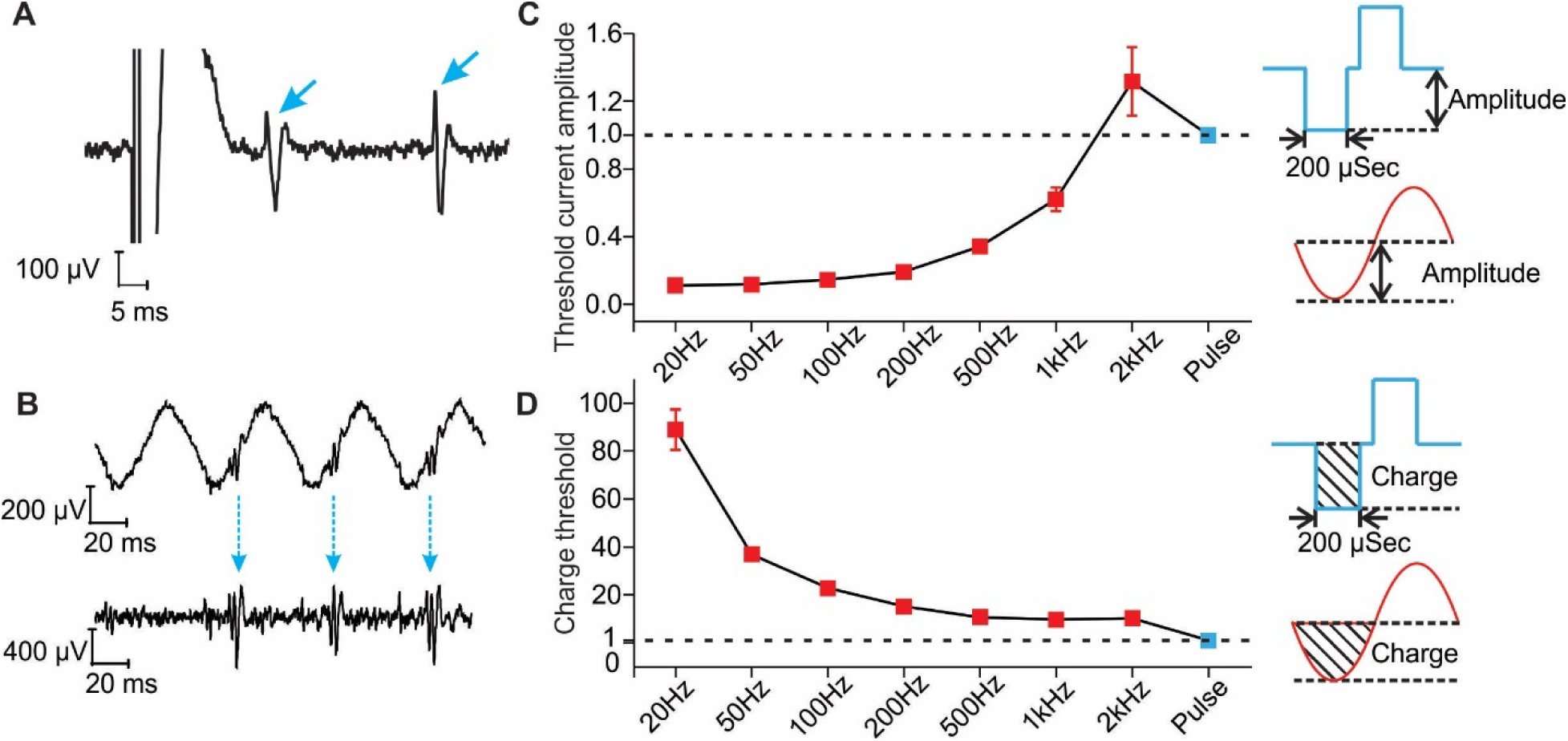
Comparison of Activation Thresholds for Sinusoidal and Pulse Stimulation. (A, B) Representative single-fiber recordings of evoked C-fiber afferents in response to pulse stimulation (200 μs) and sinusoidal stimulation (20 Hz), respectively. APs are indicated by arrows. (C) Normalized threshold amplitudes for sinusoidal stimulation across different frequencies. Threshold amplitudes are normalized to the threshold amplitude of the 200-μs pulse stimulation (dashed line). (D) Normalized charge thresholds during the cathodic period across different frequencies. Charge thresholds are normalized to the charge threshold of the 200-μs pulse stimulation (dashed line). Error bars represent the standard error of the mean (n = 10).

**Fig. 5C** displays the threshold amplitudes of sinusoidal stimulations, normalized to the threshold amplitude of the 200 μSec pulse stimulation. The threshold for 2 kHz sinusoidal stimulation is comparable to that of the pulse stimulation but over 10 times higher than the threshold for 20 Hz sinusoidal stimulation. A progressive increase in threshold was observed as the sinusoidal frequency increased from sub-kilohertz (≤100 Hz) to kilohertz ranges.

We also calculated the electric charge required to reach threshold for both pulse and sinusoidal stimulations by integrating the stimulus current over the cathodic period, i.e., the ‘charge threshold’ as illustrated in the inset of **Fig. 5D**. The charge thresholds for sinusoidal stimulation, normalized to that of the pulse stimulation, are summarized in **Fig. 5D**. While lower-frequency sinusoidal stimulations (20 and 50 Hz) have lower amplitude thresholds, they require substantially more charge to activate C-fiber axons compared to higher-frequency stimulations (1 and 2 kHz).

### Selective transmission blockage of C-fiber axons by low-frequency sinusoidal Stimulation

We previously reported that low-frequency (20 to 100 Hz) suprathreshold pulse stimulation of the DRG and peripheral nerves can reversibly block action potential transmission in C-fiber axons (Chen et al., 2022;Zhang et al., 2024). Here, we investigated whether low-frequency sinusoidal stimulation could also selectively block C-fiber axons using the ex vivo single-fiber recordings and neuromodulation protocols described in **Fig. 2**. Trains of 1.5-second sinusoidal stimulations were applied at E-Stim 2, with 0.5-second windows before and after stimulation to assess subtle changes in AP transmission initiated at E-Stim 1. **Fig. 6A** shows typical single-fiber recordings at the onset and after 10 seconds of sinusoidal stimulation. To eliminate the stimulus artifact from the sinusoidal signal, the raw recordings in **Fig. 6A** were band-pass filtered and displayed in **Fig. 6B**. Spikes were sorted using principal component analysis, with different spike clusters labeled in distinct colors. In this example, the slow-conducting C-fiber (green, indicated by arrows) was blocked after 10 seconds of 20 Hz sinusoidal stimulation (open arrowheads), while the fast-conducting A-fibers (red) and Aδ-fiber (orange) remained unblocked. When the local anesthetic lidocaine (1%) was applied to the nerve trunk, all action potential spikes were blocked, leaving only the pulse stimulus artifact, as shown in **Fig. 6C**.

**Figure 6.**
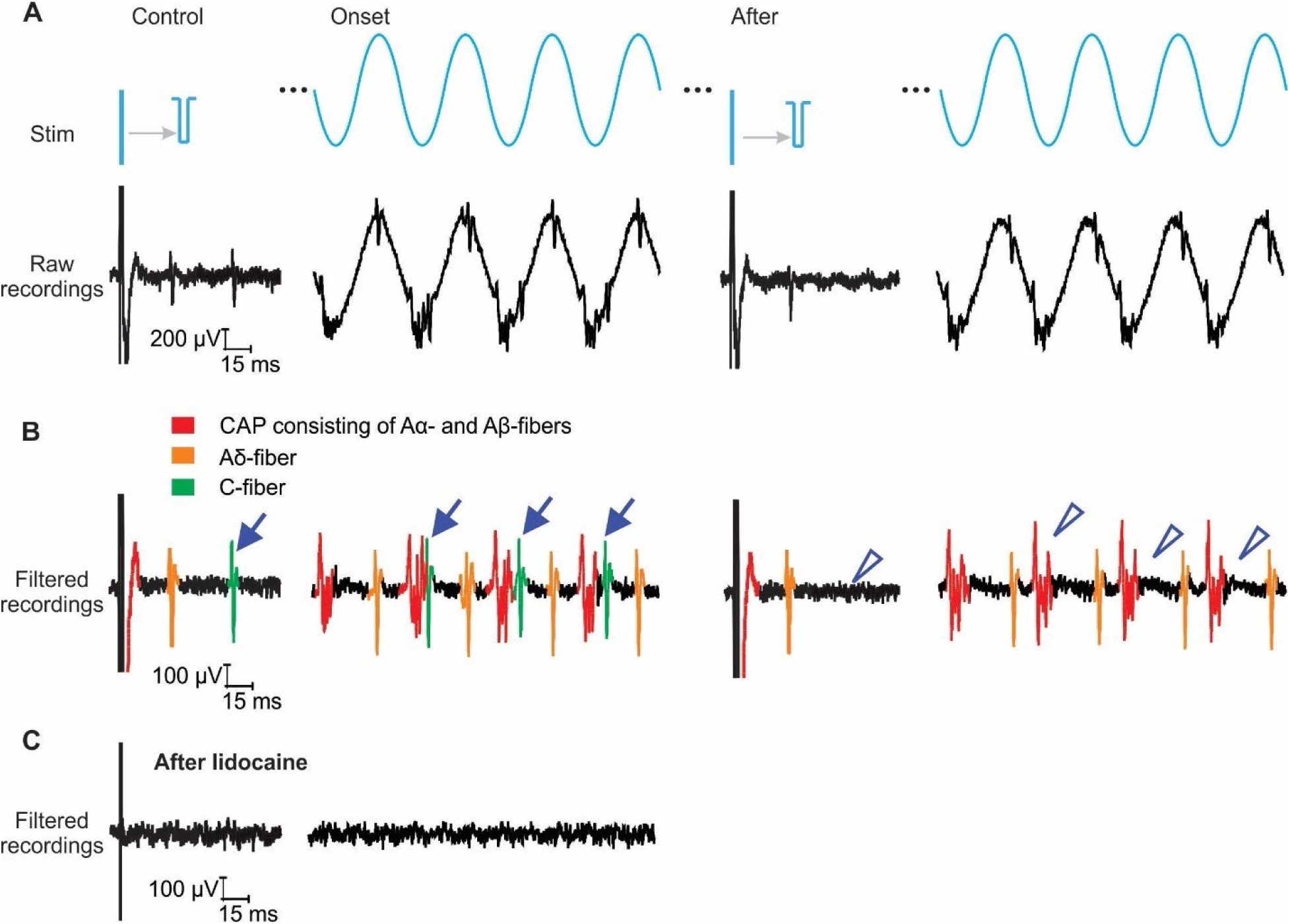
Selective Block of C-Fiber Transmission by Sinusoidal Stimulation. (A) Stimulation protocol illustrating pulse stimulation (E-Stim 1) and sinusoidal modulation (E-Stim 2). (B) Raw recording data of nerve activity during the stimulation protocol. The compound action potential (CAP) consists of signals from multiple fast-conducting Aα- and Aβ-fibers, while single-fiber recordings were isolated from Aδ- and C-fibers. (C) Filtered data demonstrating selective blockage of C-fiber transmission (green) without affecting fast A-fiber (red) or Aδ-fiber (orange) activity after 2 seconds of 20 Hz sinusoidal stimulation. (D) Complete suppression of neural activity following the application of 1% lidocaine to the nerve trunk.

We focused on evaluating the effect of low-frequency sinusoidal stimulation in blocking C-fiber afferents. The amplitude of the sinusoidal stimulation was set to 150% of the threshold required to evoke action potentials at 50 Hz; at this amplitude, the stimulation remains sub-threshold for kilohertz frequencies, as shown in **Fig. 5C**. Using the sinusoidal neuromodulation protocol described in **Fig. 2**, we applied three frequencies (20, 50, and 2000 Hz) at E-Stim 2 to assess their effect on action potential transmission evoked by pulse stimulation at E-Stim 1, measured before, during, and after sinusoidal stimulation. Representative single-fiber recordings are shown in **Fig. 7A**, illustrating a reversible block of C-fiber transmission induced by 20 Hz sinusoidal stimulation, as indicated by the open arrowheads. **Fig. 7B** summarizes the data from 11 C-fiber and 8 Aδ-fiber axons. Consistent with our previous findings using pulse stimulation (Chen et al., 2022;Zhang et al., 2024), 20 Hz sinusoidal stimulation efficiently blocked a significantly higher proportion of C-fiber axons than Aδ-fiber axons (Fisher’s exact test, p < 0.001). The blocking efficacy was reduced at 50 Hz for both C- and Aδ-fibers with no significant difference in the proportions of blocked fibers (p = 0.38). At 2 kHz, the sinusoidal stimulation remained sub-threshold and did not block any tested C- or Aδ-fibers. These results indicate that low-frequency sinusoidal stimulation, especially at 20 Hz, enables selective blockage of C-fiber axons while sparing fast-conducting A-fiber axons.

**Figure 7.**
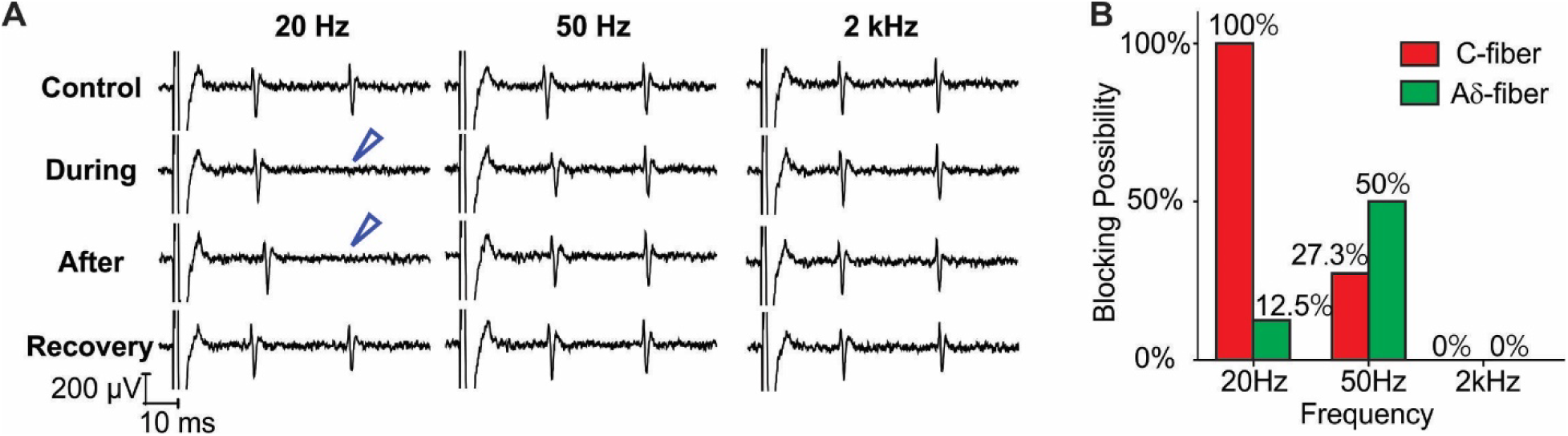
Reversible Block of C-Fiber Transmission by Low-Frequency Sinusoidal Stimulation. The stimulus amplitude was set to 150% of the threshold amplitude for 50 Hz sinusoidal stimulation. (A) Representative single-fiber recordings from peripheral nerve axons during sinusoidal neuromodulation at 20, 50, and 2000 Hz. Open arrowheads indicate instances of blocked C-fiber transmission at 20 Hz. (B) Blocking probability of C-fiber and Aδ-fiber axons at 20, 50, and 2000 Hz sinusoidal stimulation. Stimulation at 20 and 50 Hz was suprathreshold, while 2000 Hz stimulation was sub-threshold.

### Sinusoidal stimulation (20Hz) of L6 DRG reversibly blocked VMR to noxious CRD

With ex vivo single-fiber recordings, we established that 20 Hz sinusoidal stimulation selectively blocks C-fiber axons, similar to the 20-Hz pulse stimulation we reported previously (Chen et al., 2022;Zhang et al., 2024). We then assessed whether 20 Hz sinusoidal stimulation could reversibly block nociception from distal colon and rectum, which is predominantly innervated by unmyelinated C-fiber afferents. As illustrated in **Fig. 3A**, we conducted whole-animal behavior assessment in urethane-anesthetized mice to measure the VMR to CRD. We exposed bilateral L6 DRG and applied sinusoidal stimulations via two flexible MEA as shown in **Fig. 3C**. The 20-Hz sinusoidal stimulation was delivered to both pairs of electrodes in each MEA with stimulus threshold set at 40% of the motor threshold (MTh) that evokes visible limb or torso muscle twitch (Chen et al., 2022). **Fig. 8A** displays typical recordings of VMR to CRD before (control), immediately after (Neuromod), and 15 min after terminating the 20-Hz sinusoidal stimulation (recovery). As shown in **Fig. 8B**, the baseline VMR to CRD recorded from the TNBS mouse group (N = 6) was significantly higher than the saline-treated control group (N = 6) (two-way ANOVA, F_1,40_ = 7.52, P < 0.01). In the saline-treated control group, the VMR to CRD was reversibly blocked by 20-Hz stimulation (**Fig. 8C**) (two-way ANOVA with repeated measures, F_2,55_ = 179.31, P < 0.001, post-hoc comparison, p < 0.001 for ctrl vs. sine stim, p > 0.05 for ctrl vs. recovery). Similarly, 20-Hz stimulation reversibly blocked the VMR to CRD in the TNBS-treated group (**Fig. 8D**) (two-way ANOVA with repeated measures, F_2,55_ = 158.26, P < 0.001, post-hoc comparison, p < 0.001 for ctrl vs. sine stim, p > 0.05 for ctrl vs. recovery). In comparison, 2 kHz sinusoidal stimulation of the same amplitude at bilateral L6 DRG was assessed in 12 mice, which did not cause any significant changes in VMR to CRD as shown in **Fig. 8E** (two-way ANOVA with repeated measures, F_2,121_ = 2.58, P = 0.0984). This suggests that kilohertz stimulation using the amplitude determined by 20 Hz stimulation is sub-threshold.

**Figure 8.**
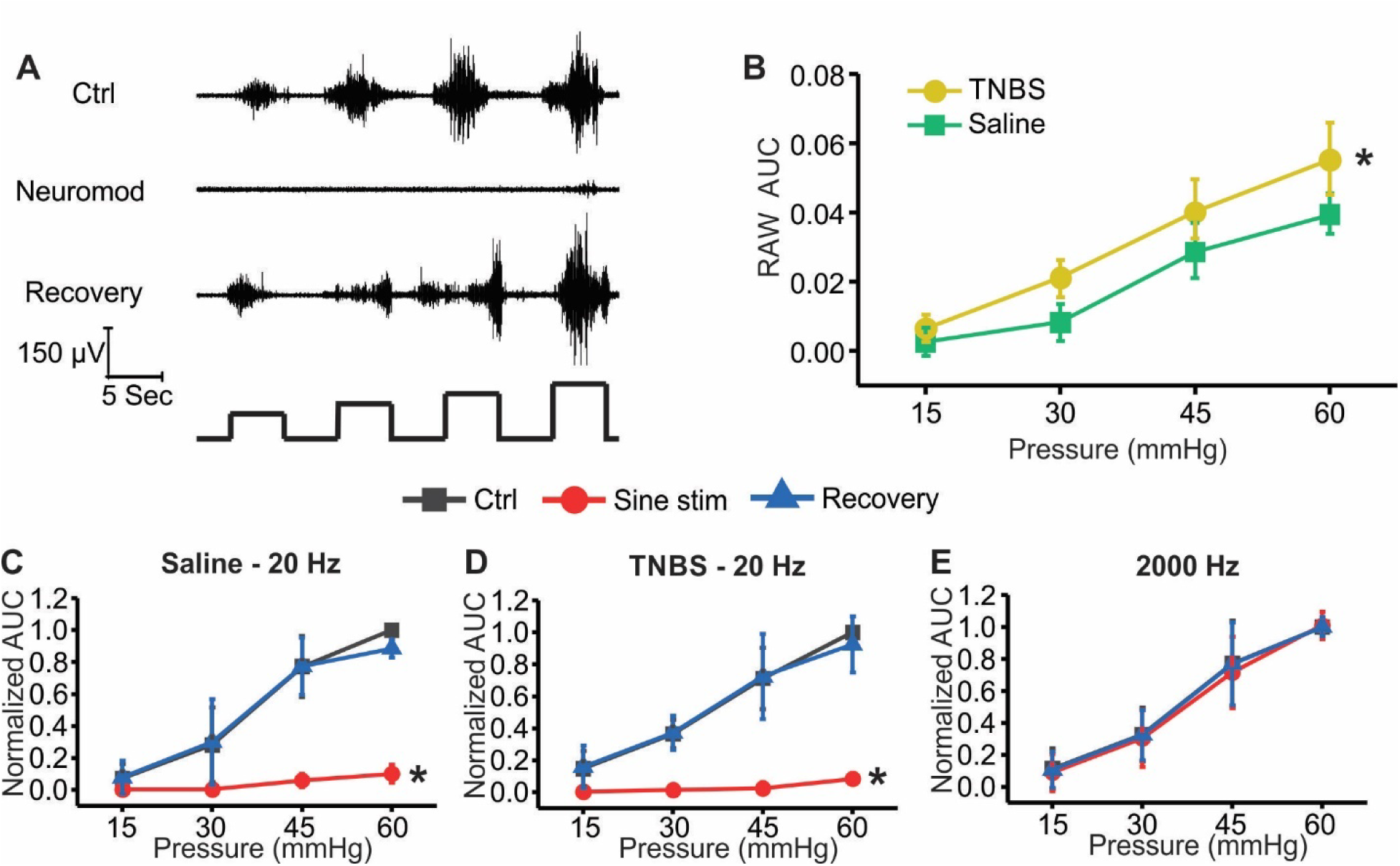
Low-frequency sinusoidal DRG stimulation (20 Hz) reversibly blocks VMR to CRD. (A) Representative EMG recordings showing VMR to CRD before (Ctrl), during (Neuromod), and after (Recovery) sinusoidal DRG stimulation. (B) Comparison of VMR to CRD between TNBS-treated and saline-treated groups. (C) 20 Hz sinusoidal DRG stimulation reversibly blocks VMR to CRD in saline-treated group. (D) Similar reversible blockade of VMR to CRD by 20 Hz stimulation in TNBS-treated group. (E) Kilohertz (2 kHz) sinusoidal stimulation at the same amplitude shows no significant effects on VMR to CRD. Data presented as mean ± SEM. * indicates significant difference (p<0.05).

### TIS of 2000 and 2020 Hz at bilateral L6 DRG reversibly blocked VMR to noxious CRD

Since TIS between two kilohertz sinusoidal waves generates a low-frequency envelope (e.g., 20 Hz) at the focal region, we further assessed the effect of TIS of L6 DRG in blocking the VMR to noxious CRD. The 2000 and 2020 Hz sinusoidal stimulations were delivered to the two electrode pairs as illustrated in **Fig. 4A**. The amplitude of the TIS stimulation was determined as 40% of the MTh for 20 Hz sinusoidal stimulation. Thus, the individual 2000 or 2020 Hz stimulation is likely sub-threshold and has no impact on the VMR to CRD as suggested by **Fig. 8E**. To vary the focal region of TIS, we delivered the following three stimulation configurations to each experiment: 1) balanced pair with both amplitudes set at 40% the MTh, 2) unbalanced pair with a 3:1 ratio, i.e., 60% and 20% the MTh, respectively, 3) unbalanced pair with a 1:3 ratio, i.e., 20% and 60% the MTh. VMR to CRD was assessed immediately after all three TIS configurations in 6 saline-treated mice, and the highest level of inhibition is summarized in **Fig. 9A**, showing reversible blockage of VMR to CRD (two-way ANOVA with repeated measures, F_2,55_ = 107.29, p<0.001, post-hoc comparison, p < 0.001 for Ctrl vs. TIS, p > 0.05 for Ctrl vs. Recovery). Similarly, TIS reversibly blocked VMR to CRD in 6 TNBS-treated mice as reported in **Fig. 9B** (two-way ANOVA with repeated measures, F_2,55_ = 94.26, p < 0.001, post-hoc comparison, p < 0.001 for Ctrl vs. TIS, p > 0.05 for Ctrl vs. Recovery). The VMR to CRD following the three TIS configurations were ranked and pooled together in **Fig. 9C** (6 saline-treated and 6 TNBS-treated), indicating significant inhibition by the first- and second-ranked TIS, but not by the third-ranked TIS (Two-way ANOVA with repeated measures, F_3,272_ = 121.75, p < 0.001, post-hoc comparison, p < 0.001 for 1st vs. Ctrl and 2nd vs. Ctrl).

**Figure 9.**
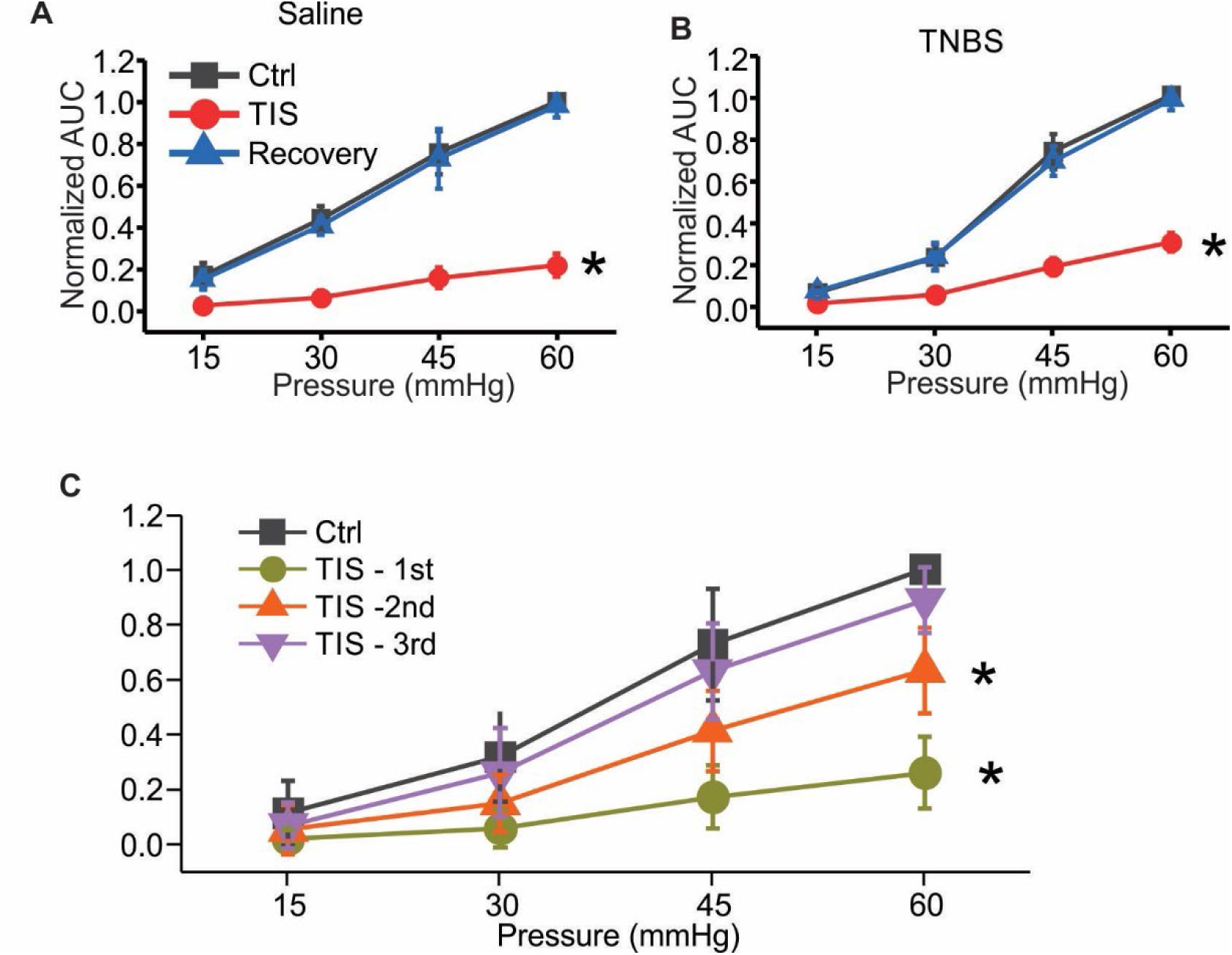
TIS effectively suppresses visceromotor responses (VMR) to colorectal distension (CRD). Each mouse was assessed with three TIS configurations using amplitude ratios of 1:3, 1:1, and 3:1. (A) Highest level of VMR suppression to CRD by one of the three TIS configurations in saline-treated control group (n = 6). (B) Highest level of VMR suppression to CRD by one of the three TIS configurations in TNBS-treated group (n = 6). (C) Ranked suppression of VMR to CRD by the three TIS configurations, pooled from both groups (n = 12). Data are presented as mean ± SEM.

### TIS of L6 DRG causes no apparent immediate immune responses

We conducted histological staining of DRG undergoing hours long TIS to evaluate the possible immediate immune-responses caused by kilohertz stimulation. We use monocyte infiltration to assess the immediate immune responses, which is quantified by antibody staining of F4/80 harvested DRG sections as shown in **Fig. 10**. No significant difference (t-test, p = 0.08) in fluorescent ratio of F4/80 over DAPI were detected between the L6 DRG (stimulated, 0.10±0.01, n = 11 in 4 mice) and L5 DRG (not stimulated, 0.11±0.01, n = 11 in 4 mice). There is also no apparent recruitment of spherical-shaped monocytes from the bloodstream, as the F4/80-stained cells exhibit an amorphous shape.

**Figure 10.**
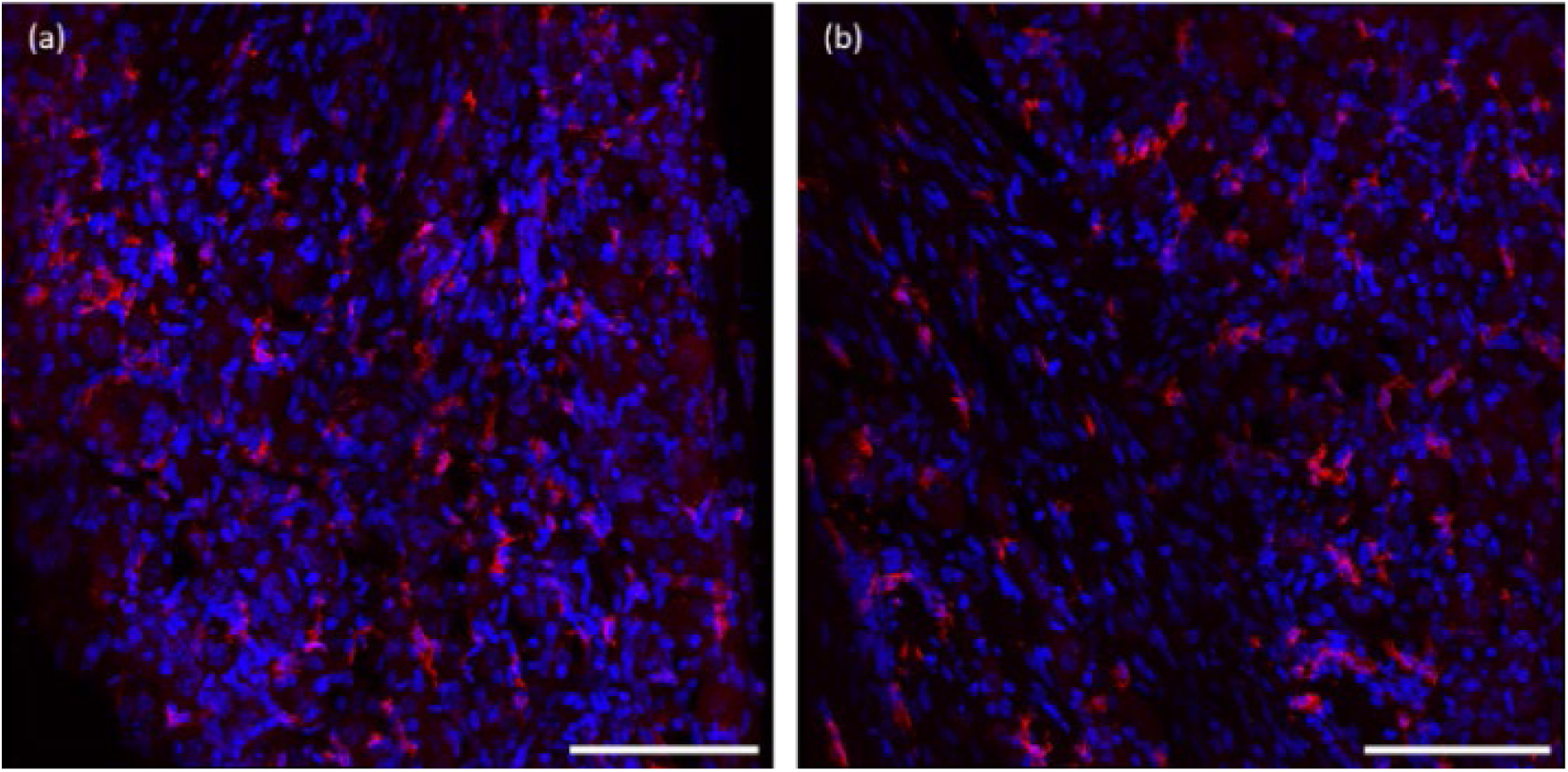
Immunostaining of F4/80-immunoreactive macrophages (red) in DRG tissue. (a) Electrically stimulated L6 DRG. (b) Control L5 DRG from the same mouse receiving no stimulation. Tissue sections were counterstained with DAPI (nuclei in blue). Scale bar: 100 μm.

## Discussion

The current study contributes to the extensive literature exploring selective C-fiber nociceptor blockade as a treatment for chronic pain. Typically occurring in the absence of apparent tissue injury, chronic pain is characterized by long-term maladaptive changes in both the central and peripheral nervous systems (CNS and PNS) (Gold and Gebhart, 2010;Kuner, 2010). Mounting evidence reveals that central sensitization in the CNS is primarily driven by enhanced spatial and temporal summation of synaptic input from peripheral C-fibers, particularly C-fiber nociceptors in deep tissues (see (Latremoliere and Woolf, 2009) for a review). In contrast to the immediate sharp sensation of acute pain, chronic pain patients often describe their pain as achy (53%), throbbing (28%), burning (22%), and/or dull (20%) (Jensen et al., 2013), indicating activation of C-fiber nociceptors. Furthermore, human microneurographic studies demonstrate that C-fiber sensitization is present in chronic pain patients experiencing hyperalgesia and allodynia (Ørstavik et al., 2003;Serra et al., 2014). This C-fiber sensitization underlies prolonged pain sensation that outlasts stimulus duration and likely serves as a major mechanism driving the persistence of chronic pain. Consequently, targeting the dorsal root ganglia (DRG) with tunable transcranial interferential stimulation (TIS) presents a promising strategy for addressing chronic pain by directly suppressing peripheral sensitization. It is hypothesized that by removing this driver of central sensitization, CNS plasticity could, over time, reverse the sensitization process, potentially allowing patients to recover from allodynia and hyperalgesia.

The current study is the first to implement TIS on dorsal root ganglia (DRG) stimulation, aiming to enhance the tunability of the modulatory area beyond the stimulus sites. TIS employs two or more stimulations at kilohertz carrier frequencies to generate interfered sinusoidal signals with a low-frequency pattern of amplitude modulation at focal regions between the stimulation sites. Notably, the focal regions of TIS can be adjusted by altering the amplitude ratio of the two interfering stimulation signals (Guo et al., 2023).TIS was proposed decades ago under various names, such as interferential current therapy and interferential current stimulation (Goats, 1990;Agharezaee and Mahnam, 2015). This technique has gained significant research interest following the seminal study by Grossman et al., which demonstrated the potential for activating deep brain tissues with noninvasive surface stimulation (Grossman et al., 2017). Their study utilized in vivo whole-cell patch-clamp recordings in mouse brain to reveal the activation of neural somata by TIS delivered from outside the skull.The initial interpretation of TIS’s neural mechanism as low-pass filtering of neural membrane (Grossman et al., 2017) was challenged by subsequent theoretical studies on the fundamental physics of TIS. These studies indicated the necessary role of ion-channel mediated transmembrane current rectification in extracting the sinusoidal envelope signal to drive neural activation (Mirzakhalili et al., 2020). Furthermore, theoretical and experimental studies suggest that the stimulus intensity of focal brain regions with TIS delivered from outside the skull is likely below the threshold of neural activation, indicating a mechanism of sub-threshold neural modulation (Rampersad et al., 2019;Lee et al., 2020;Howell and McIntyre, 2021).Theoretical analysis also indicates that high-intensity kilohertz stimulation can potentially cause axonal conduction block near the stimulation site (Grossman et al., 2017;Budde et al., 2023). However, this effect has yet to be validated by direct experimental measurements on blocked neuronal tissues. Despite these unanswered questions, several clinical studies are currently assessing the neuromodulatory effect of implementing TIS in stimulating deep brain structures (Collavini et al., 2021;Ma et al., 2021;Zhu et al., 2022;Violante et al., 2023;Vassiliadis et al., 2024).

The aforementioned uncertainties associated with TIS largely stem from the scarcity of experimental data directly measuring the evoked neural activities. The continuous kilohertz stimulation introduces substantial stimulus artifacts in extracellular recordings, hindering direct measurement of evoked neural spikes even in in vitro neural culture systems (Ahtiainen et al., 2024). To address this challenge, we employed GCaMP6s recordings from whole mouse DRG to directly assess the activation of afferent neurons by TIS. This optical recording approach, which we recently established, is unaffected by electrical stimulus artifacts (Guo et al., 2019;Bian et al., 2021). We also utilized a flexible and transparent microelectrode array (MEA) to cover the contour of the DRG surface, enabling concurrent electrical stimulation and optical recordings of the DRG neurons underneath. This novel approach was recently reported by our group (Ryu et al., 2024). Our optical recordings from individual DRG neurons revealed that: 1) DRG neurons are efficiently activated by 20-Hz sinusoidal current stimulation; 2) kilohertz sinusoidal stimulation (2000, 2020 Hz) at the same amplitude is subthreshold and did not activate any neurons, and 3) TIS enables activation of DRG neurons away from the stimulation sites and steering of activated regions by adjusting the amplitude ratio of the two kilohertz stimulus signals. Compared to the complex application of TIS in the brain, our data indicate that TIS produces a clear and steerable activation zone away from the two interfering stimulation sites in the DRG. This effect is likely due to the relatively homogeneous neuron types in the DRG (i.e., the lack of inhibitory neurons), absence of a sophisticated neural network, and small geometry of the DRG, with the focal TIS region only sub-millimeters away from the stimulation sites.

The envelope of the TIS can be extracted by neurons as a rectified low-frequency sinusoidal wave (Mirzakhalili et al., 2020), which justifies the current study’s focus on low-frequency sinusoidal stimulation in activating nerve axons. In contrast to pulse waveforms, only a few studies have investigated continuous sinusoidal waves in stimulating neurons and axons (Koga et al., 2005;Sundar and González-Cueto, 2006). While the relation between stimulus pulse width and activation threshold has been well established as the strength-duration curve for electrical pulse stimulation (Geddes and Bourland, 1985), a similar characterization with sinusoidal stimulations at different frequencies has not been reported in the literature. In this study, we conducted single-fiber recordings from split sciatic nerve axons in an ex vivo preparation (Chen et al., 2017b;Chen et al., 2018;Guo et al., 2020), enabling accurate measurement of the activation threshold of C-fiber axons as a function of sinusoidal stimulation frequency, as shown in **Fig. 5C**. The progressive increase in activation threshold with increased sinusoidal frequency (or reduced period) is consistent with the strength-duration curve for pulse stimulation, where the threshold also increases with reduced pulse width. Our results indicate that the threshold to activate C-fiber sciatic nerve axons is much higher at kilohertz frequencies than at 20 Hz, which aligns with a prior experimental study showing more efficient activation of C-fiber axons with 5-Hz sinusoidal stimulation compared to 250 and 2000 Hz (Koga et al., 2005). The drawback of using sinusoidal stimulation to activate neurons over pulse stimulation is evident: as shown in **Fig. 5D**, nearly 100 times more electrical charge is required with a 20 Hz sine-wave stimulation than a pulse stimulation of 0.2 mSec duration.

This study also reports the first nerve blocking experiments with sinusoidal stimulation, demonstrating the blocking effect on Aδ- and C-fiber axons with 20 to 50 Hz stimulation. We recently conducted systematic studies using electrical pulse stimulation and reported that low-frequency pulse stimulation (10 to 100 Hz) effectively blocks Aδ- and C-fiber afferents with stimulus delivered either at the DRG (Chen et al., 2022) or peripheral nerve trunk (Zhang et al., 2024). In contrast, fast-conducting Aα- and Aβ-fiber afferents are not blocked by this subkilohertz range of stimulation (Patel and Butera, 2015;Patel and Butera, 2018). By implementing a synchronized stimulation protocol, we were the first to demonstrate that DRG stimulation causes progressive activity-dependent conduction slowing until transmission block, which gradually recovers 5 to 10 min after terminating the DRG stimulation (Chen et al., 2022). This activity-dependent conduction slowing was previously reported in microneurographic studies on human C-fiber sensory axons, where a 2-Hz stimulation protocol was implemented to cause significant conduction slowing in putative C-fiber nociceptors (e.g., (Serra et al., 1999;Serra et al., 2004)). Consistent with prior pulse studies (Chen et al., 2022;Zhang et al., 2024), suprathreshold sinusoidal stimulation appears necessary for causing conduction blockage. In the absence of evoked action potential spikes, subthreshold sinusoidal stimulation at a wide range of frequencies (10 to 1000 Hz) did not affect afferent transmission, as indicated by the unchanged afferent conduction delay before and after stimulation. Our recent computational simulation study suggests that this conduction block with activity-dependent slowing is likely contributed to by dysregulated intra-axonal sodium and potassium concentrations (Zhang et al., 2024). This provides a mechanistic interpretation for the propensity of low-frequency stimulation (10 – 100Hz) to block small-diameter C- and Aδ-fibers over large-diameter fibers.

We assessed the efficacy of DRG stimulation with TIS by conducting behavioral visceromotor response (VMR) to colorectal distension (CRD), a well-established and quantitative assessment of visceral pain in mouse models (Christianson and Gebhart, 2007). To accommodate the invasive surgeries, such as exposing the bilateral L6 DRG, we recently established an anesthesia protocol that enables robust and repeatable measurement of VMR to CRD for hours in mice under urethane anesthesia (Zhang et al., 2023). We implemented intracolonic enema of TNBS to induce behavioral visceral hypersensitivity following our previously reported methods (Feng et al., 2012b;Chen et al., 2022;Zhang et al., 2023), and demonstrated that DRG stimulation with TIS reversibly blocked the VMR to CRD in both control and TNBS groups. Notably, we showed that by varying the amplitude ratio of the two interfering stimulus signals, the blocking effect of VMR to CRD can be finely adjusted. This is consistent with the GCaMP6 recordings showing that the focal region of activation by TIS can be steered by varying the amplitude ratio. Thus, the current study provides proof of concept that TIS enables tunable neuromodulation of DRG neurons to effectively block visceral nociception arising from sensitized C-fiber nociceptors.

Regarding the safety of TIS, we demonstrated that DRG neurons can be efficiently activated by TIS with stimulating signal amplitudes below those used for 200-µSec pulse stimulation (**Fig. 5C**). Our results with F4/80 staining showed the absence of monocyte infiltration in the DRG following hours-long TIS, indicating no early immune responses in the DRG. This contrasts with the detection of mild colonic inflammation by the same F4/80 staining protocol following the enema of zymosan (Feng et al., 2012a).

We focused on DRG neuromodulation with TIS as a proof of concept for treating chronic visceral pain, despite the widespread implementation of spinal cord stimulation (SCS) and dorsal peripheral nerve stimulation (dPNS) in managing chronic pain. This choice was largely due to DRG stimulation harnessing the mechanism of afferent transmission block, which is not easily achieved by SCS or dPNS. Unlike DRG stimulation, SCS and dPNS employ electrodes in contact with epineural nerve trunks. Epineural nerve stimulation typically activates large, myelinated A-fiber axons at a much lower stimulus threshold than small unmyelinated C-fiber axons (Zhang et al., 2024). C-fiber axons are generally not activated with clinical settings of implantable neurostimulators. In our recent ex vivo experimental recordings, we had to use a glass suction electrode to sufficiently evoke C-fiber axons in peripheral nerves by creating a high-impedance seal between the electrode and surrounding bath solution (Zhang et al., 2024). Consequently, SCS and dPNS usually do not target unmyelinated C-fiber afferents for treating pain. Instead, they suppress pain by activating low-threshold A-fiber afferents to evoke paresthesia, which inhibits nociceptive signaling in the spinal cord. However, this approach is not applicable to managing visceral pain, as visceral organs generally lack prominent innervation of A-fiber afferents for evoking paresthesia. Moreover, engaging the central inhibitory neural circuit with A-fiber activation only temporarily suppresses pain and does not remove heightened C-fiber peripheral drive, which can exacerbate central sensitization over time. Sensitized C-fiber nociceptors are generally unaffected by SCS or dPNS and continue to assault the CNS, driving further maladaptive changes (Latremoliere and Woolf, 2009). Therefore, targeting the lumbosacral DRG with electrical stimulation can be a promising strategy for treating patients with prolonged visceral hypersensitivity and pain by selectively blocking C-fiber nociception.

## Conclusions

This study presents a novel approach to chronic pain management by implementing TIS on DRG to selectively block C-fiber nociceptors. Our findings demonstrate that TIS enables tunable neuromodulation of DRG neurons, effectively blocking visceral nociception arising from sensitized C-fiber nociceptors. The study provides compelling evidence through GCaMP6 recordings, single-fiber recordings, and behavioral assessments that TIS can produce a clear and steerable activation zone in the DRG, with the ability to fine-tune the blocking effect by adjusting the amplitude ratio of interfering stimulus signals. Unlike traditional methods such as SCS and dPNS, TIS on DRG targets unmyelinated C-fiber afferents directly, addressing a critical gap in current pain management strategies, particularly for visceral pain. The low stimulation amplitude required and the absence of early immune responses in the DRG suggest a favorable safety profile. These findings pave the way for developing more targeted and effective treatments for chronic visceral pain, potentially offering relief to patients suffering from this debilitating condition.

## Acknowledgments

This work was supported by NINDS U01 NS113873, NIDDK R01 DK120824, and NSF 1844762 grants awarded to BF.

## References

1. Agharezaee, M., and Mahnam, A. (2015). A computational study to evaluate the activation pattern of nerve fibers in response to interferential currents stimulation. Medical & biological engineering & computing 53, 713–720.

2. Ahtiainen, A., Leydolph, L., Tanskanen, J.M., Hunold, A., Haueisen, J., and Hyttinen, J.A. (2024). Electric field temporal interference stimulation of neurons in vitro. Lab on a Chip 24, 3945–3957.

3. Anand, P., Aziz, Q., Willert, R., and Van Oudenhove, L. (2007). Peripheral and central mechanisms of visceral sensitization in man. Neurogastroenterology & Motility 19, 29–46.

4. Bian, Z., Guo, T., Jiang, S., Chen, L., Liu, J., Zheng, G., and Feng, B. (2021). High-Throughput Functional Characterization of Visceral Afferents by Optical Recordings From Thoracolumbar and Lumbosacral Dorsal Root Ganglia. Frontiers in Neuroscience 15.

5. Budde, R.B., Williams, M.T., and Irazoqui, P.P. (2023). Temporal interference current stimulation in peripheral nerves is not driven by envelope extraction. Journal of neural engineering 20, 026041.

6. Chao, D., Mecca, C.M., Yu, G., Segel, I., Gold, M.S., Hogan, Q.H., and Pan, B. (2021). Dorsal root ganglion stimulation of injured sensory neurons in rats rapidly eliminates their spontaneous activity and relieves spontaneous pain. PAIN 162.

7. Chao, D., Zhang, Z., Mecca, C.M., Hogan, Q.H., and Pan, B. (2020). Analgesic dorsal root ganglionic field stimulation blocks conduction of afferent impulse trains selectively in nociceptive sensory afferents. PAIN 161.

8. Chen, L., Guo, T., Siri, S., and Feng, B. (Year). “Frequency-dependent modulation of action potential transmission in afferent axons by DRG stimulation”, in: Neuroscience Meeting Planner, Chicago, IL: Society for Neuroscience 2019, Oct. 19 – 23, Chicago, IL.).

9. Chen, L., Guo, T., Siri, S., and Feng, B. (Year). “Suprathreshold DRG stimulation blocks action potential transmission in afferent axons while subthreshold stimulation does not”, in: Proceedings of the Biomedical Engineering Society 2019 Annual Fall Meeting, Oct. 12 – 16, Philadelphia, PA.).

10. Chen, L., Guo, T., Zhang, S., Smith, P.P., and Feng, B. (2022). Blocking peripheral drive from colorectal afferents by subkilohertz dorsal root ganglion stimulation. PAIN 163, 665–681.

11. Chen, L., Ilham, S.J., and Feng, B. (2017a). Pharmacological Approach for Managing Pain in Irritable Bowel Syndrome: A Review Article. Anesth Pain Med 7, e42747.

12. Chen, L., Ilham, S.J., Guo, T., Emadi, S., and Feng, B. (2017b). In vitro multichannel single-unit recordings of action potentials from mouse sciatic nerve. Biomed Phys Eng Express 3, 045020.

13. Chen, L., Lucas, R.F., and Feng, B. (2018). A Novel System to Measure Infants’ Nutritive Sucking During Breastfeeding: the Breastfeeding Diagnostic Device (BDD). IEEE J Transl Eng Health Med 6, 2700208.

14. Christianson, J.A., and Gebhart, G.F. (2007). Assessment of colon sensitivity by luminal distension in mice. Nature Protocols 2, 2624–2631.

15. Collavini, S., Fernández-Corazza, M., Oddo, S., Princich, J.P., Kochen, S., and Muravchik, C.H. (2021). Improvements on spatial coverage and focality of deep brain stimulation in pre-surgical epilepsy mapping. Journal of Neural Engineering 18, 046004.

16. Dijk, G., Ruigrok, H.J., and O’connor, R.P. (2021). PEDOT: PSS-coated stimulation electrodes attenuate irreversible electrochemical events and reduce cell electropermeabilization. Advanced Materials Interfaces 8, 2100214.

17. Feng, B., and Gebhart, G.F. (2011). Characterization of silent afferents in the pelvic and splanchnic innervations of the mouse colorectum. Am J Physiol Gastrointest Liver Physiol 300, G170–180.

18. Feng, B., and Guo, T. (2020). Visceral pain from colon and rectum: the mechanotransduction and biomechanics. Journal of Neural Transmission 127, 415–429.

19. Feng, B., La, J.H., Schwartz, E.S., Tanaka, T., Mcmurray, T.P., and Gebhart, G.F. (2012a). Long-term sensitization of mechanosensitive and -insensitive afferents in mice with persistent colorectal hypersensitivity. Am J Physiol Gastrointest Liver Physiol 302, G676–683.

20. Feng, B., La, J.H., Tanaka, T., Schwartz, E.S., Mcmurray, T.P., and Gebhart, G.F. (2012b). Altered colorectal afferent function associated with TNBS-induced visceral hypersensitivity in mice. Am J Physiol Gastrointest Liver Physiol 303, G817–824.

21. Geddes, L., and Bourland, J. (1985). The strength-duration curve. IEEE transactions on biomedical engineering, 458–459.

22. Goats, G.C. (1990). Interferential current therapy. British Journal of Sports Medicine 24, 87–92.

23. Gold, M.S., and Gebhart, G.F. (2010). Nociceptor sensitization in pain pathogenesis. Nat Med 16, 1248–1257.

24. Grossman, N., Bono, D., Dedic, N., Kodandaramaiah, S.B., Rudenko, A., Suk, H.-J., Cassara, A.M., Neufeld, E., Kuster, N., Tsai, L.-H., Pascual-Leone, A., and Boyden, E.S. (2017). Noninvasive Deep Brain Stimulation via Temporally Interfering Electric Fields. Cell 169, 1029–1041.e1016.

25. Günter, C., Delbeke, J., and Ortiz-Catalan, M. (2019). Safety of long-term electrical peripheral nerve stimulation: review of the state of the art. Journal of neuroengineering and rehabilitation 16, 1–16.

26. Guo, T., Bian, Z., Trocki, K., Chen, L., Zheng, G., and Feng, B. (2019). Optical recording reveals topological distribution of functionally classified colorectal afferent neurons in intact lumbosacral DRG. Physiological Reports 7, e14097.

27. Guo, T., Chen, L., Tran, K., Ghelich, P., Guo, Y.-S., Nolta, N., Emadi, S., Han, M., and Feng, B. (2020). Extracellular single-unit recordings from peripheral nerve axons in vitro by a novel multichannel microelectrode array. Sensors and Actuators B: Chemical 315, 128111.

28. Guo, T., Patel, S., Shah, D., Chi, L., Emadi, S., Pierce, D.M., Han, M., Brumovsky, P.R., and Feng, B. (2021). Optical clearing reveals TNBS-induced morphological changes of VGLUT2-positive nerve fibers in mouse colorectum. American Journal of Physiology-Gastrointestinal and Liver Physiology 320, G644–G657.

29. Guo, W., He, Y., Zhang, W., Sun, Y., Wang, J., Liu, S., and Ming, D. (2023). A novel non-invasive brain stimulation technique:“Temporally interfering electrical stimulation”. Frontiers in Neuroscience 17, 1092539.

30. Harrison, C., Epton, S., Bojanic, S., Green, A.L., and Fitzgerald, J.J. (2018). The efficacy and safety of dorsal root ganglion stimulation as a treatment for neuropathic pain: a literature review. Neuromodulation: Technology at the Neural Interface 21, 225–233.

31. Howell, B., and Mcintyre, C.C. (2021). Feasibility of interferential and pulsed transcranial electrical stimulation for neuromodulation at the human scale. Neuromodulation: Technology at the Neural Interface 24, 843–853.

32. Hulisz, D. (2004). The burden of illness of irritable bowel syndrome: current challenges and hope for the future. J Manag.Care Pharm. 10, 299–309.

33. Jänig, W. (1996). Neurobiology of visceral afferent neurons: neuroanatomy, functions, organ regulations and sensations. Biological psychology 42, 29–51.

34. Jensen, M.P., Johnson, L.E., Gertz, K.J., Galer, B.S., and Gammaitoni, A.R. (2013). The words patients use to describe chronic pain: Implications for measuring pain quality. PAIN® 154, 2722–2728.

35. Knotkova, H., Hamani, C., Sivanesan, E., Le Beuffe, M.F.E., Moon, J.Y., Cohen, S.P., and Huntoon, M.A. (2021). Neuromodulation for chronic pain. The Lancet 397, 2111–2124.

36. Koga, K., Furue, H., Rashid, M.H., Takaki, A., Katafuchi, T., and Yoshimura, M. (2005). Selective Activation of Primary Afferent Fibers Evaluated by Sine-Wave Electrical Stimulation. Molecular Pain 1, 1744–8069-1741-1713.

37. Kuner, R. (2010). Central mechanisms of pathological pain. Nat Med 16, 1258–1266.

38. Latremoliere, A., and Woolf, C.J. (2009). Central Sensitization: A Generator of Pain Hypersensitivity by Central Neural Plasticity. The Journal of Pain 10, 895–926.

39. Lee, S., Lee, C., Park, J., and Im, C.-H. (2020). Individually customized transcranial temporal interference stimulation for focused modulation of deep brain structures: a simulation study with different head models. Scientific reports 10, 11730.

40. Liang, M.-Z., Yang, J.-W., Chen, H., Cheng, M.-Y., and Chen, L. (2023). Quantitative monitoring of neuronal regeneration by functional assay and wireless neural recording. bioRxiv, 2023.2009. 2021.558793.

41. Linderoth, B., and Foreman, R.D. (1999). Physiology of spinal cord stimulation: review and update. Neuromudulation: Technology at the Neural Interface 2, 150–164.

42. Lovell, R.M., and Ford, A.C. (2012). Effect of gender on prevalence of irritable bowel syndrome in the community: systematic review and meta-analysis. Am J Gastroenterol 107, 991–1000.

43. Ma, R., Xia, X., Zhang, W., Lu, Z., Wu, Q., Cui, J., Song, H., and Fan, C. (2021). “High gamma and beta temporal interference stimulation in the human motor cortex improves motor functions. Front. Neurosci., 15, 800436”.).

44. Melzack, R., and Wall, P.D. (1965). Pain mechanisms: a new theory. Science 150, 971–979.

45. Mirzakhalili, E., Barra, B., Capogrosso, M., and Lempka, S.F. (2020). Biophysics of temporal interference stimulation. Cell Systems 11, 557–572. e555.

46. Ørstavik, K., Weidner, C., Schmidt, R., Schmelz, M., Hilliges, M., Jørum, E., Handwerker, H., and Torebjörk, E. (2003). Pathological C-fibres in patients with a chronic painful condition. Brain 126, 567–578.

47. Patel, Y.A., and Butera, R.J. (2015). Differential fiber-specific block of nerve conduction in mammalian peripheral nerves using kilohertz electrical stimulation. J Neurophysiol 113, 3923–3929.

48. Patel, Y.A., and Butera, R.J. (2018). Challenges associated with nerve conduction block using kilohertz electrical stimulation. Journal of neural engineering 15, 031002.

49. Rampersad, S., Roig-Solvas, B., Yarossi, M., Kulkarni, P.P., Santarnecchi, E., Dorval, A.D., and Brooks, D.H. (2019). Prospects for transcranial temporal interference stimulation in humans: a computational study. NeuroImage 202, 116124.

50. Ryu, J., Qiang, Y., Chen, L., Li, G., Han, X., Woon, E., Bai, T., Qi, Y., Zhang, S., and Liou, J.Y. (2024). Multifunctional Nanomesh Enables Cellular-Resolution, Elastic Neuroelectronics. Advanced Materials, 2403141.

51. Serra, J., Campero, M., Bostock, H., and Ochoa, J. (2004). Two types of C nociceptors in human skin and their behavior in areas of capsaicin-induced secondary hyperalgesia. J Neurophysiol 91, 2770–2781.

52. Serra, J., Campero, M., Ochoa, J., and Bostock, H. (1999). Activity-dependent slowing of conduction differentiates functional subtypes of C fibres innervating human skin. J Physiol 515 (Pt 3), 799–811.

53. Serra, J., Collado, A., Solà, R., Antonelli, F., Torres, X., Salgueiro, M., Quiles, C., and Bostock, H. (2014). Hyperexcitable C nociceptors in fibromyalgia. Annals of Neurology 75, 196–208.

54. Siegel, S., Noblett, K., Mangel, J., Bennett, J., Griebling, T.L., Sutherland, S.E., Bird, E.T., Comiter, C., Culkin, D., and Zylstra, S. (2018). Five-year followup results of a prospective, multicenter study of patients with overactive bladder treated with sacral neuromodulation. The Journal of urology 199, 229–236.

55. Sundar, S., and González-Cueto, J.A. (Year). “On the activation threshold of nerve fibers using sinusoidal electrical stimulation”, in: 2006 International Conference of the IEEE Engineering in Medicine and Biology Society: IEEE), 2908-2911.

56. Sunshine, M.D., Cassarà, A.M., Neufeld, E., Grossman, N., Mareci, T.H., Otto, K.J., Boyden, E.S., and Fuller, D.D. (2021). Restoration of breathing after opioid overdose and spinal cord injury using temporal interference stimulation. Communications Biology 4, 107.

57. Vassiliadis, P., Stiennon, E., Windel, F., Wessel, M.J., Beanato, E., and Hummel, F.C. (2024). Safety, tolerability and blinding efficiency of non-invasive deep transcranial temporal interference stimulation: first experience from more than 250 sessions. Journal of Neural Engineering 21, 024001.

58. Verne, G.N., Robinson, M.E., Vase, L., and Price, D.D. (2003). Reversal of visceral and cutaneous hyperalgesia by local rectal anesthesia in irritable bowel syndrome (IBS) patients. Pain 105, 223–230.

59. Verne, G.N., Sen, A., and Price, D.D. (2005). Intrarectal lidocaine is an effective treatment for abdominal pain associated with diarrhea-predominant irritable bowel syndrome. J Pain 6, 493–496.

60. Verrills, P., Mitchell, B., Vivian, D., Cusack, W., and Kramer, J. (2019). Dorsal Root Ganglion Stimulation Is Paresthesia-Independent: A Retrospective Study. Neuromodulation: Technology at the Neural Interface 22, 937–942.

61. Verrills, P., Vivian, D., Mitchell, B., and Barnard, A. (2011). Peripheral nerve field stimulation for chronic pain: 100 cases and review of the literature. Pain medicine 12, 1395–1405.

62. Violante, I.R., Alania, K., Cassarà, A.M., Neufeld, E., Acerbo, E., Carron, R., Williamson, A., Kurtin, D.L., Rhodes, E., and Hampshire, A. (2023). Non-invasive temporal interference electrical stimulation of the human hippocampus. Nature neuroscience 26, 1994–2004.

63. Zhang, S., Chen, L., and Feng, B. (2023). An anesthesia protocol for robust and repeatable measurement of behavioral visceromotor responses to colorectal distension in mice. Front Pain Res (Lausanne) 4, 1202590.

64. Zhang, S., Chen, L., Ladez, S.R., Seferge, A., Liu, J., and Feng, B. (2024). Blocking Aδ-and C-fiber neural transmission by sub-kilohertz peripheral nerve stimulation. Frontiers in Neuroscience 18, 1404903.

65. Zhang, T.C., Janik, J.J., and Grill, W.M. (2014). Mechanisms and models of spinal cord stimulation for the treatment of neuropathic pain. Brain Res 1569, 19–31.

66. Zhou, Q., Price, D.D., and Verne, G.N. (2008). Reversal of visceral and somatic hypersensitivity in a subset of hypersensitive rats by intracolonic lidocaine. Pain 139, 218–224.

67. Zhu, Z., Xiong, Y., Chen, Y., Jiang, Y., Qian, Z., Lu, J., Liu, Y., and Zhuang, J. (2022). Temporal interference (TI) stimulation boosts functional connectivity in human motor cortex: a comparison study with transcranial direct current stimulation (tDCS). Neural plasticity 2022, 7605046.

68. Zia, J.K., Lenhart, A., Yang, P.-L., Heitkemper, M.M., Baker, J., Keefer, L., Saps, M., Cuff, C., Hungria, G., and Videlock, E.J. (2022). Risk factors for abdominal pain–related disorders of gut–brain interaction in adults and children: A systematic review. Gastroenterology 163, 995–1023. e1023

